# Chemoproteomic profiling of *Plasmodium falciparum* Hsp90 inhibition reveals functional link to DNA replication pathways

**DOI:** 10.64898/2026.08.28.747854

**Authors:** Gaini Ibrasheva, Yueqi Chen, Michael E. Chirgwin, Caitlyn J. Hughes, Michael C. Fitzgerald, Emily R. Derbyshire

## Abstract

*Plasmodium falciparum* heat shock protein 90 (PfHsp90) is a promising antimalarial target, but the molecular pathways influenced by its inhibition remain poorly understood. Herein, we leveraged chemoproteomic profiling employing geldanamycin and XL888 Hsp90 inhibitors to investigate proteins and pathways dependent on the chaperone during the *Plasmodium* blood stage. This study revealed 131 proteins reduced in abundance after inhibition, of which 40% co-immunoprecipitated with PfHsp90. Bioinformatic analyses identified DNA replication as the most enriched pathway. This link was investigated in phenotypic studies demonstrating reduced parasite DNA content after PfHsp90 inhibition. To assess nascent DNA synthesis, we utilized a 7-deaza-7-ethynyl-2’-deoxyadenosine (EdA) assay, yielding dual-stage attenuation of nucleoside incorporation following Hsp90 inhibition. We further show that parasite co-treatment with Hsp90 and DNA replication inhibitors produces synergistic interactions, highlighting the therapeutic potential of the discovered link. Overall, these findings expand our understanding of PfHsp90 function and uncover novel PfHsp90-dependent pathways.

## INTRODUCTION

Malaria, a devastating infectious disease caused by intracellular *Plasmodium* parasites, remains a global threat with over 282 million cases and 610,000 deaths in 2024.^1^ Among the five species that infect humans, *Plasmodium falciparum* accounts for over 90% of mortalities. Despite ongoing eradication efforts, growing resistance to first-line artemisinin-based combination therapies (ACTs) exacerbates the challenge of malaria control, emphasizing the need to identify novel parasite drug targets and partner drug combinations.^2–4^

*Plasmodium* parasites undergo multiple morphological changes as they replicate in a mosquito vector and human host. During the asexual blood stage, *Plasmodium* progresses through ring, trophozoite, and schizont stages, culminating in the red blood cell (RBC) rupture and release of 16 – 32 newly replicated, infective merozoites into the host bloodstream, which gives rise to malaria’s clinical symptoms.^5,6^ To survive throughout their lifecycle, the parasites must withstand drastic environmental changes, including heat shock as they travel from a cold-blooded mosquito to a warm-blooded mammal, where cyclical fever can fluctuate between 37 °C to 42 °C.^7,8^ *Plasmodium*’s ability to acclimate to such alternating environments is key to its survival and necessitates heat shock proteins (Hsps) to provide a first line defense against proteotoxic stress.

Heat shock protein 90 (Hsp90) is a conserved molecular chaperone that plays a central role in maintaining cellular proteostasis.^9^ By undergoing ATP-driven conformational cycles, Hsp90 supports the folding, maturation, and regulation of a specific set of proteins, termed clients.^10^ The *P. falciparum* Hsp90 (PfHsp90) has been widely recognized as a compelling antimalarial target due to its continuous expression throughout the parasite’s lifecycle,^11–13^ hypersensitivity of its ATPase activity to inhibition,^14^ and structural and functional differences from the human Hsp90 homolog.^15,16^ Previous proteomic analyses of Hsp90 in model systems indicate that the chaperone supports 10 – 20% of eukaryotic proteomes,^17,18^ with clients implicated in diverse pathways, including signal transduction,^19,20^ cell cycle regulation,^21,22^ apoptosis,^23,24^ cell growth,^25^ and translation.^26^ However, the client repertoire of Hsp90 is divergent across different species. While PfHsp90 clients are predicted to be involved in critical parasite pathways, including chromatin remodeling, protein translation, and drug resistance;^27,28^ the physical PfHsp90 interactome remains largely unresolved with only two clients validated to date.^29,30^

Hsp90 is one of the most highly expressed proteins in eukaryotes and is essential across all *P. falciparum* life stages. This has challenged efforts to genetically alter the protein to gain deeper insights into its function.^11,31^ On the other hand, proteomic profiling of Hsp90 inhibition has expanded our understanding of Hsp90-regulated processes in various human cancer cell lines,^32–35^ and more recently, in the protozoan parasite *Leishmania mexicana*.^36^ In *P. falciparum*, the PfHsp90 interactome has been investigated in one study using a pulldown approach and by our group via mapping protein thermal stability changes upon inhibitor treatment.^13,28^ These efforts provided valuable insights into PfHsp90 biology, linking the chaperone to the chloroquine resistance transporter and the proteasome, respectively; however as these studies were completed using parasite lysate, there is a need to interrogate chaperone function within the native cellular context. Given Hsp90’s transient, low-affinity interactions, which are influenced by physiological conditions and co-chaperones, disrupting Hsp90 within intact parasites would enable a more comprehensive mapping of its functional pathways.^37,38^

In this study, we performed proteome-wide profiling of PfHsp90 in-cell inhibition to obtain insights into proteins and biological processes dependent on the chaperone. To identify conserved proteome responses while minimizing chemotype-specific off-target effects, we employed two distinct PfHsp90 inhibitors, geldanamycin and XL888. Our analysis identified 131 proteins whose abundances were reduced by both inhibitors, ∼40% of which physically associated with PfHsp90 based on a co-immunoprecipitation (co-IP) analysis. Bioinformatic analyses yielded a strong enrichment in DNA replication, suggesting a putative functional link. Further investigation revealed impaired late-stage parasite development and reinvasion, as well as reduced nucleoside incorporation following Hsp90 inhibitor treatments. Promisingly, we found that PfHsp90 disruption sensitizes *P. falciparum* to DNA replication inhibitors, providing therapeutic relevance of the connection. Together, this work demonstrates the importance of PfHsp90 in supporting DNA replication and further reveals over a hundred potential interactors, including those unique to *Plasmodium,* for future investigation.

## RESULTS

### Geldanamycin and XL888 target PfHsp90 in parasites

To interrogate the PfHsp90-dependent proteome, we implemented an approach whereby PfHsp90 was inhibited in live *P. falciparum* blood stage parasites prior to bottom-up proteomic analysis. For this study, two Hsp90 inhibitors, the benzoquinone ansamycin geldanamycin (GA) and the tropane-derived XL888 (XL), were selected for their well-characterized Hsp90 inhibition profiles and previous application as Hsp90 probes.^35,39,40^ (**Figure 1A**). GA is a well-established PfHsp90 inhibitor, while XL, which advanced to phase Ib/II clinical trials for targeting human Hsp90 (HsHsp90) in combination therapy for solid tumors, was recently identified as a PfHsp90-binding molecule.^13,41^ GA and XL bind to purified PfHsp90 (K_i_=2.3 nM for GA and K_i_=27 nM for XL) with 3-4-fold greater affinity than HsHsp90, and GA associates with PfHsp90 in parasite lysate based on immobilization-mediated pulldown assays.^12–14^ We sought to identify proteome-level responses in the parasite shared by both molecules, thereby minimizing the contribution of compound-specific off-target effects. We further reasoned that their use at parasite EC_50_ concentrations in erythrocytes that lack nuclei, ribosomes and other major organelles, should minimize confounding effects from HsHsp90 inhibition.^42^

**Fig 1.**
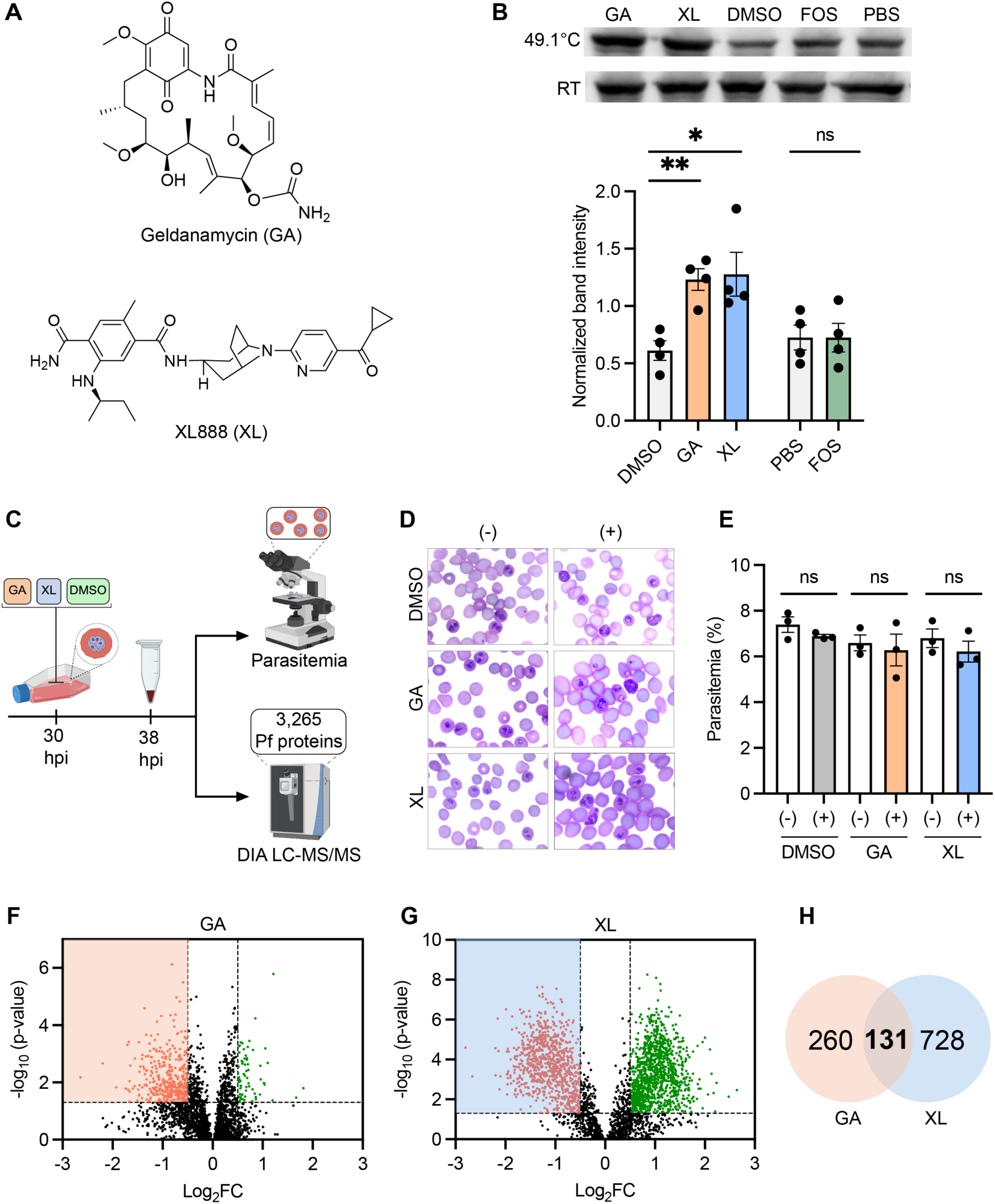
PfHsp90-dependent proteins identified via global proteomic analysis. (A) Structures of geldanamycin (GA) and XL888 (XL). (B) Representative Western blot detecting PfHsp90 from *P. falciparum* lysate treated with 100 µM GA, XL, or the negative control fosmidomycin (FOS) at room temperature (RT) or 49.1 °C. The vehicle controls were at 5% (GA and XL: DMSO and FOS: PBS). PfHsp90 band intensities at 49.1 °C were normalized to respective bands at RT. Total input was normalized with Bradford dye and confirmed with Ponceau S stain. Data shown as means ± SEM, *n*=4 biological replicates; not significant (ns) >0.05, *<0.05, **<0.01; Welch’s t-test. (C) Schematic representation of parasite treatment and analysis. (D-E) *P. falciparum*-infected RBCs before (-) and after (+) treatment with 0.3 µM GA or XL, or 0.5% DMSO at 30 hpi for 8 hours. (D) Representative light microscopy images of smears. (E) Quantification of parasitemia. Data shown as means ± SEM, *n*=3 biological replicates, analyzing >900 RBCs for each condition; not significant (ns) >0.05; Welch’s t-test. (F and G) The -log_10_ (Bonferroni corrected ANOVA *p*-value) is plotted against Log_2_(fold change compound/DMSO) for GA (F) and XL (G). Dashed line on *X*-axis indicates the Log_2_FC >|0.5| cutoff and dashed line on *Y*-axis indicates the *p*-value <0.05 cutoff. Proteins that significantly increased (green dots) or decreased (red dots) abundance following treatment are indicated. *n*=3 biological replicates. (H) Venn diagram highlighting the overlap of significantly reduced proteins between GA and XL treatments.

To first validate the physical interaction of GA and XL with PfHsp90 in lysate, we employed a thermal shift assay (TSA) coupled with Western blot, where ligand binding stabilizes the protein against heat-induced denaturation, leading to an increased abundance of soluble protein. Similar to previous studies,^30,43^ we employed an HsHsp90 antibody that detects PfHsp90 due to their significant sequence homology (64% identity).^44^ The HsHsp90 antibody was first validated with cellular lysate and purified protein, where a strong signal was detected in *P. falciparum* lysate and purified PfHsp90, but not in RBCs (**Figure S1A**). This validated the detection of PfHsp90 in enriched parasite lysate, whereas HsHsp90 in RBCs was below the detection limit, supporting the use of the antibody for TSA. A previous study utilizing thermal proteome profiling (TPP) in *P. falciparum* lysate demonstrated BX2819 (Hsp90 inhibitor) binding-induced stabilization of PfHsp90 at 50.6 °C.^13^ Building on this, the temperature range for GA and XL was initially profiled between 47.1–53.1 °C. For this, *P. falciparum* isolated from host RBCs were lysed and treated with 100 µM compound for 1 hour. Lysates were then split and subjected to thermal challenge, where we observed ligand-induced stabilization between 49.1–51.8 °C with the greatest detected stabilization at 49.1 °C when compared to DMSO vehicle control (**Figure S1B**). Following this preliminary study, the experiment was repeated at 49.1 °C with inclusion of a negative control fosmidomycin (FOS), an antimalarial that targets the non-mevalonate pathway of isoprenoid synthesis and is therefore unrelated to Hsp90 (**Figure 1B**).^45^ DMSO was the vehicle control for GA and XL while FOS was dissolved in PBS, consistent with the supplier recommendation and prior studies that formulated FOS in aqueous buffers.^46–48^ Compound- and vehicle-treated lysate was then split, where one half was incubated at room temperature (RT) and the other was heated to 49.1 °C. The anti-Hsp90 Western blot band intensities of soluble PfHsp90 recovered after heat treatment were quantified and normalized to their respective bands at RT. We observed a significant increase in the PfHsp90 band with GA and XL at 49.1 °C (*p* <0.05) but detected no significant change with FOS treatment (*p*=0.995). Together, these studies support the binding of GA and XL to Hsp90 in *P. falciparum*.

### PfHsp90 inhibition alters protein abundances in the parasite proteome

We next sought to employ GA and XL in a global proteomic analysis to identify overall changes induced by PfHsp90 inhibition and provide insights into potential PfHsp90-dependent proteins. First, we aimed to identify suitable drug concentrations that would engage PfHsp90 without significant parasite death within 24–36 hours post invasion (hpi), when protein synthesis is most active.^49,50^ *P. falciparum*-infected RBCs at 30 hpi were incubated with GA or XL at their EC_50_ concentrations (0.3 µM)^12,13^ or 0.5% DMSO for 8 hours (**Figure 1C**). To assess growth, parasite blood smears were taken prior to and after compound treatments for parasitemia (parasite content in blood) quantification. The smears demonstrated no notable changes in parasite morphology or parasitemia with any treatments (**Figures 1D** and **1E**). These compound concentrations and incubation time were identified as suitable for subsequent proteomic analysis.

*P*. *falciparum*-infected RBCs were treated with GA and XL at their EC_50_ concentration for 8 hours at 30 hpi. Parasites were then isolated from RBCs, lysed and analyzed by LC-MS/MS on an Orbitrap Astral with a data-independent acquisition (DIA) bottom-up approach (*n*=3 biological replicates). A total of 3,265 parasite proteins were detected, achieving >60% of the predicted 5,361 *P. falciparum* proteome (UniProt ID: UP000001450) and representing one of the highest coverages reported to date (**Figure 1C, Table S1**).^51,52^ Proteins exhibiting Log_2_FC of >0.5 or <- 0.5 with *p*-value<0.05 were considered significantly altered, revealing hundreds of proteins increased and decreased in abundance after GA or XL treatment (**Figures 1F** and **1G, Table S2A**). Generally, fewer changes were observed following GA treatment, likely owing to its known slow *k*_on_ rate and limited bioavailability.^53–55^ In GA-treated parasites, the number of proteins with decreased abundance was almost 6-fold greater than those with increased abundance, consistent with the targeting of a molecular chaperone and client destabilization (**Figure 1F**). On the other hand, XL-treated parasites exhibited 1.3-fold more proteins with increased versus decreased abundance, suggesting greater off-target effects or a compensatory stress response (**Figure 1G**). Importantly, an overlap of 131 significantly reduced proteins between both compounds was observed, highlighting scaffold-independent proteome-level responses (**Figure 1H**). In contrast, only 33 overlapping proteins with increased abundance were noted, and a preliminary functional analysis on these yielded no substantial enrichment in any pathway, suggesting they may be false positives. Since our primary objective was to identify proteins that depend on PfHsp90 for proper folding and function, we focused on the 131 overlapping proteins with reduced abundance. Collectively, these constitute a prioritized high confidence set of PfHsp90-dependent proteins, making them strong candidates for further investigation.

### Global proteomic analysis yields conserved and *Plasmodium*-specific candidate PfHsp90 interactors

To evaluate our 131 overlapping proteins more broadly, we sought to leverage the substantial body of work done in model systems, where yeast represents the most extensively characterized Hsp90 interactome.^13,56,57^ To compare our findings to these prior studies, we examined if our high confidence parasite proteins have human or yeast orthologs (**Figure 2A, Table S2B)**. Among the 131 proteins, 32% had *Homo sapiens* orthologs and 27% had *Saccharomyces cerevisiae* orthologs. These protein subsets were mined for established Hsp90 interactors, where 43% of the human orthologs and 72% of the yeast orthologs had physical or genetic evidence of Hsp90 interaction. To next connect the two independent analyses, the human interactors were mapped to the yeast genome, which identified 14 proteins with orthologs (**Table S2C)**. Among these, 11 were identified in our study, suggesting a conserved interaction between human, yeast, and *Plasmodium*. Notably, 7 of the 11 conserved proteins (Q8IC16, Q8IDF0, Q8I2N2, Q8I3A1, Q7KQM1, Q8IM38, and Q8IKF0) are involved in DNA replication, mitotic cell cycle, and ribonucleotide biosynthesis. The remaining 4 (Q7KQL5, Q8IAU5, O77344, and Q8I566) are associated with ubiquitination, microtubule formation, signal transduction, and folate metabolism. The overlap between our findings and prior reports in model organisms supports the validity of our experimental approach.

**Fig 2.**
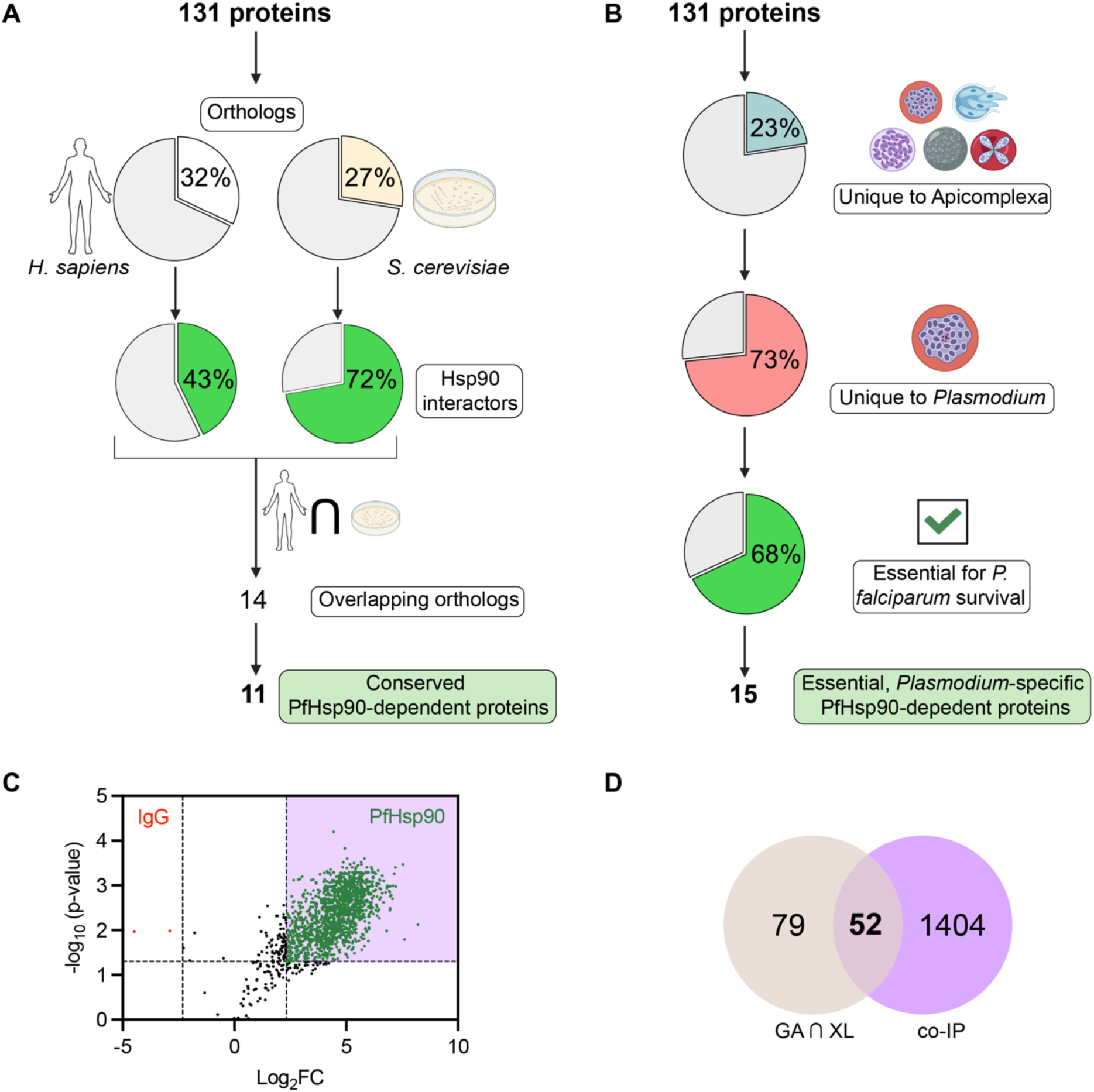
Validation of proteomic approach via ortholog analysis and co-IP. (A) *H. sapiens* and *S. cerevisiae* orthologs of the 131 high confidence PfHsp90-dependent proteins. Orthologs were analyzed for previously reported physical and genetic interactions with Hsp90 isoforms, identifying 11 in all 3 species. (B) Proteins unique to Apicomplexa (30) or *Plasmodium* (22) were identified among the PfHsp90-dependent proteins, where 15 are predicted to be essential based on a genome-wide mutagenesis screen.^58^ (C) The -log_10_ (Benjamin Hochberg FDR corrected *p*-value) is plotted against Log_2_(fold change PfHsp90/IgG). Dashed line on *X*-axis indicates FC >|5| cutoff and dashed line on *Y*-axis indicates the *p*-value <0.05 cutoff. Proteins significantly enriched in PfHsp90 (green dots) or IgG (red dots) are indicated. (D) Venn diagram highlighting the overlap in proteins between GA & XL inhibition-based and co-IP proteomic analyses.

We observed that most of our 131 high confidence proteins lacked orthologs in yeast and human model systems, which was expected as almost two-thirds of *P. falciparum* proteins are unique to this organism due to its evolutionary divergence and highly AT-rich genome.^59^ Beyond this, a third of the *Plasmodium* proteome remains unannotated, further complicating functional predictions.^60^ PfHsp90-dependent proteins that are unique and essential to *Plasmodium* are compelling for further investigation due to their potential for selective targeting. Therefore, we evaluated if the reduced proteins were *Plasmodium*-specific and assessed their indispensability for parasite survival based on a previous genome-wide mutagenesis screen, which allows for the identification of essential genes even when they have unknown function.^58^ We identified 23% of the 131 proteins as unique to the Apicomplexa phylum where almost three fourths were present exclusively in *Plasmodium* (**Figure 2B, Table S2D**). These proteins were then cross-referenced with the mutagenesis screen, yielding 15 PfHsp90-dependent *Plasmodium*-unique proteins expected to be essential for parasite survival. Notably, almost all had unknown or putative functions, representing novel candidate drug targets for selective inhibition.

Next, we sought to assess whether our PfHsp90-dependent proteins physically interact with the chaperone using a proteomic PfHsp90 co-IP profiling. Briefly, lysates isolated from *P. falciparum*-infected RBCs were incubated with A/G beads coupled with either an anti-Hsp90 antibody or an anti-IgG antibody as a non-specific control. To minimize disruptions to transient interactions, parasites were immediately processed for co-IP after isolation. Eluted proteins were analyzed by bottom-up DIA to support detection of low abundance interactors. A fold change of >5 and a *p*-value<0.05 were selected as enrichment thresholds, leading to the identification of 1,456 proteins as PfHsp90-enriched (**Figure 2C**). Importantly, only 2 proteins were IgG-enriched, demonstrating the specificity of the approach. When this co-IP analysis was compared to the inhibition-based proteomics, an overlap of 52 proteins (40%) was noted (**Figure 2D, Table S2E**). Further, >50% of the conserved and >30% of the essential *Plasmodium*-specific proteins (**Figures 2A-B)** were enriched by PfHsp90 co-IP (**Table S2F**). These proteins represent candidate PfHsp90 interactors likely dependent on its function.

### PfHsp90-dependent proteins are enriched in DNA replication and nuclear localization

To investigate potential functional interactions among the reduced proteins, a protein-protein interaction (PPI) network was generated (**Figure 3A**). The cluster analysis revealed a strong enrichment in DNA-templated DNA replication with 68 genes mapping to this pathway. Gene Ontology (GO) analysis of Biological Process (BP) and Cellular Component (CC) was performed to further characterize enriched functional pathways (**Figures 3B** and **3C**). The BP results indicated a substantial enrichment in DNA replication and DNA maintenance, and most proteins were predicted to localize to the nucleus based on the CC analysis. Beyond DNA replication, we also noted an enrichment in DNA repair and associated pathways in GO BP, aligning with previous literature (**Figure 3B**).^29,61^ GO CC analysis further revealed significant enrichment of replication fork, nuclear chromosome, and replisome components (**Figure 3C**).

**Fig 3.**
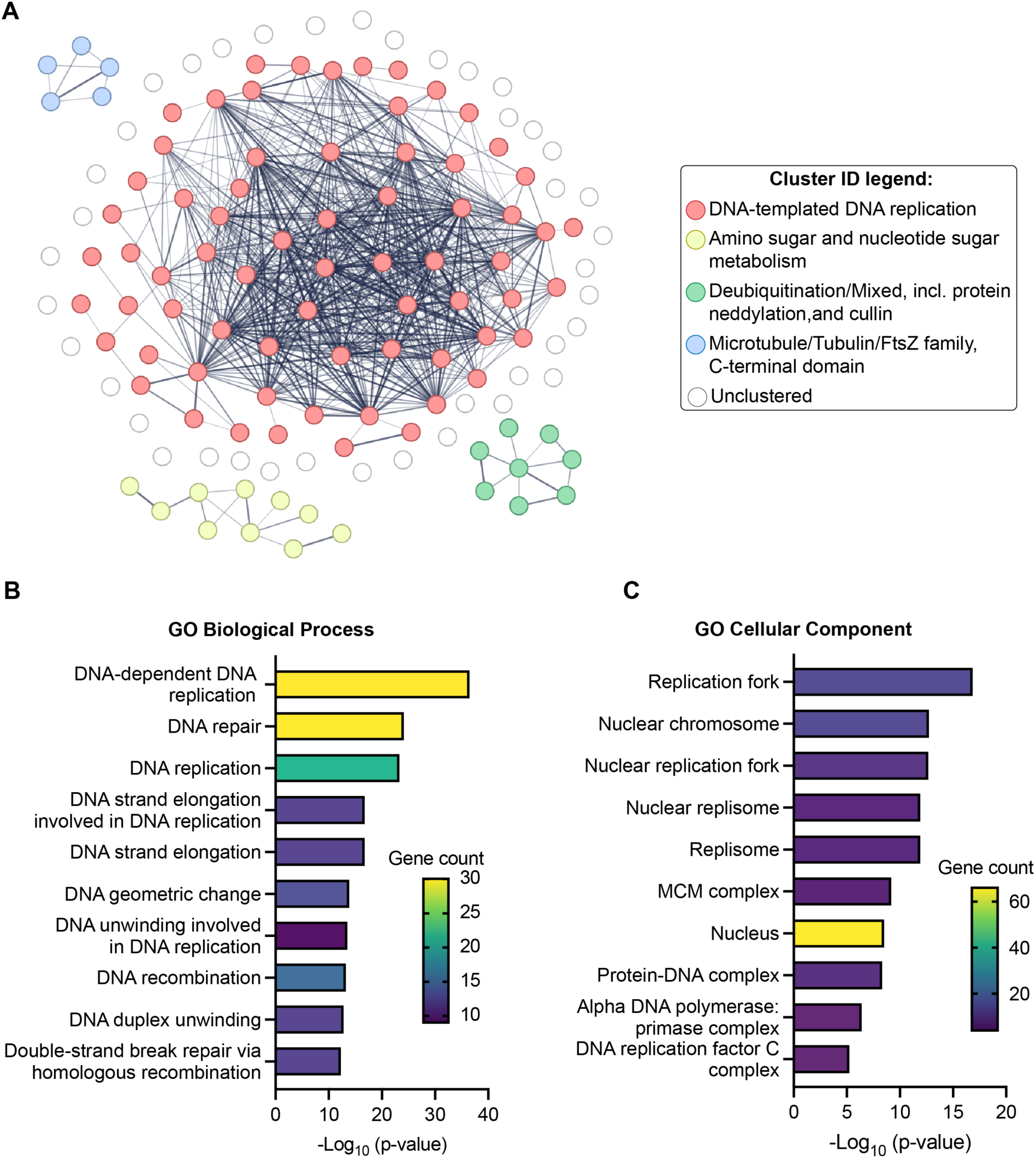
Computational analysis to inform function of PfHsp90-dependent proteins. (A) Interaction network of PfHsp90-dependent proteins, where nodes represent proteins, the color corresponds to the cluster ID, and the edge line thickness indicates the strength of the interaction prediction. (B and C) Gene Ontology (GO) enrichment analysis demonstrating the top 10 most significantly enriched GO terms (*p* <0.05, Fisher’s exact test) in Biological Process (B) and Cellular Component (C). *X*-axis represents -Log_10_ *p-*value while the color of the bars and placement on the *Y*-axis correspond to the number of genes associated with the GO term.

Due to incomplete functional annotation of *Plasmodium* proteins, only 80 and 94 proteins out of 131 PfHsp90-dependent proteins were analyzed in GO BP and CC, respectively. To predict the associations of the excluded proteins, a functional analysis of their structural homologs (Dali) was performed,^62^ resulting in 43% of proteins predicted to be involved in DNA replication and associated pathways **(Figure S2A, Table S3A)**. Proteins excluded from the GO CC were examined for their likely subcellular localization (DeepLoc 2.0),^63^ yielding up to 62% of proteins predicted to have a nuclear localization signal **(Figures S2B, Table S3B)**.

Given this computational evidence that DNA replication was impacted by Hsp90 inhibition, we examined if the 30 PfHsp90-dependent proteins involved in DNA-dependent DNA replication (**Figure 2B**) were within the co-IP dataset. We found that two-thirds (20) of these were pulled down by PfHsp90, including multiple subunits of DNA polymerases α, δ, and ε, DNA primase, MCM2-7 complex, as well as replication protein A1 and replication factor C (**Table S3C**). Taken together, these data present compelling evidence for a functional and physical link between PfHsp90 and DNA replication machinery.

### Hsp90 inhibition attenuates total and nascent DNA during the *P. falciparum* blood stage

We next sought to test our hypothesis that PfHsp90 is linked to parasite DNA replication. First, changes in stage distribution following GA and XL treatments from the trophozoite-to-late trophozoite stage (30–38 hpi) or trophozoite-to-schizont stage (30–42 hpi) were assessed (**Figure 4A**). As expected, we observed a marked increase in schizonts and rings in the vehicle DMSO control at 38 and 42 hpi, consistent with successful schizogony and reinvasion. This progression was impacted by the positive control E64d, a late-stage cysteine protease inhibitor known to impact parasite development, at its EC_50_ concentration (0.6 µM, *n*=3). E64d caused an accumulation of schizonts at 38 hpi, followed by a partial increase in rings at 42 hpi. At the 10xEC_50_ concentration of E64d, no increase in rings was observed at 42 hpi (**Figure S3A**). E64d also led to swollen food vacuoles, aligning with previous reports and supporting the robustness of this assay in detecting established stage-specific phenotypes.^64,65^ In contrast, GA and XL treatments at their EC_50_ and 10xEC_50_ concentrations led to persistence of trophozoites, fewer schizonts and no increase in rings. These observations suggest impaired schizogony and a failure to reinvade with Hsp90 inhibitor treatment.

**Fig 4.**
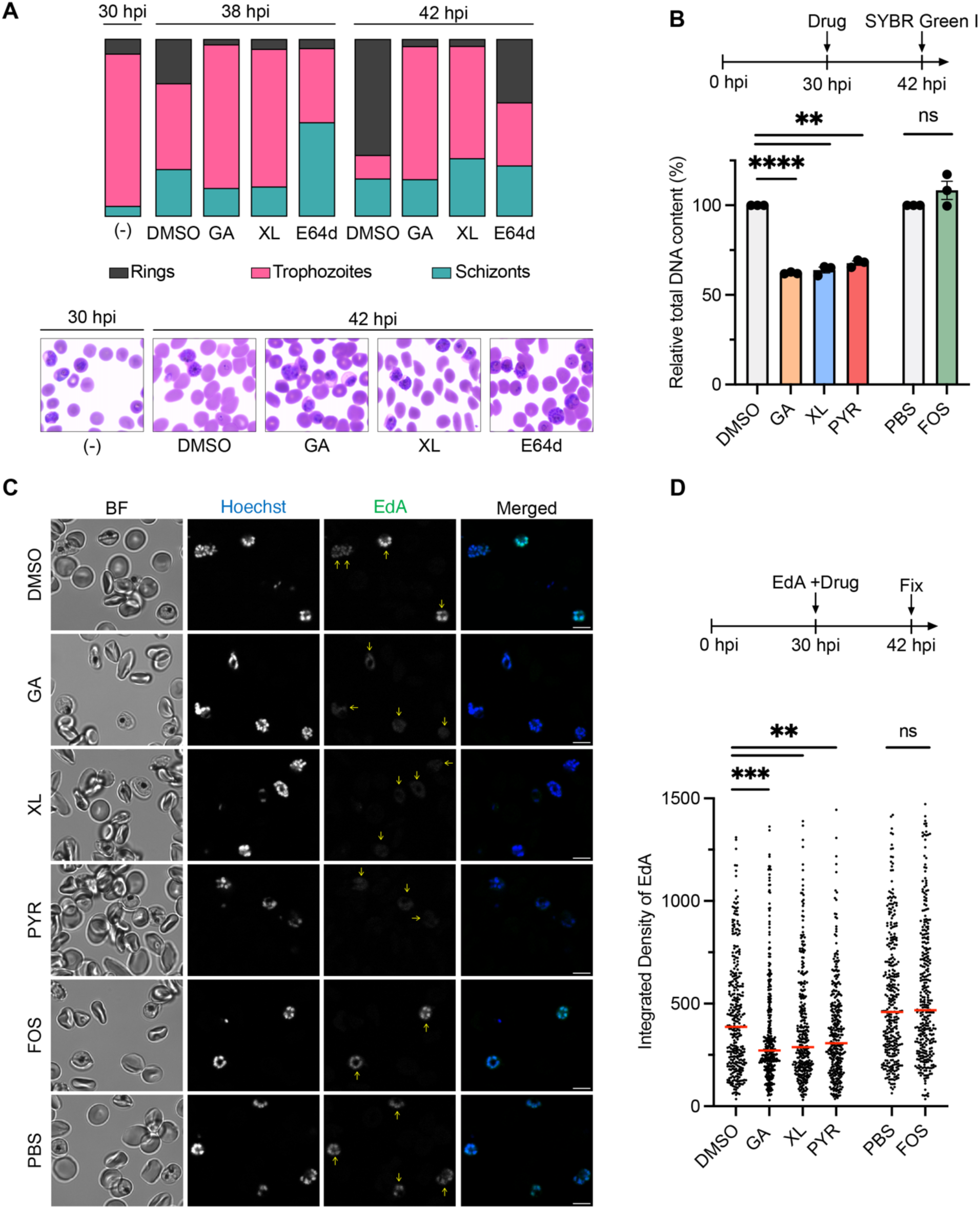
PfHsp90 inhibition reduces DNA abundance and nucleoside incorporation. (A) *P. falciparum*-infected RBCs were treated with GA, XL, and E64d at EC_50_ concentrations for 8 or 12 hours at 30 hpi. DMSO was at 0.5%. Stage progression was assessed via light microscopy analysis of smears. *n*=3 biological replicates analyzing >290 parasites for each condition. (B) *P. falciparum*-infected RBCs were treated with GA, XL, PYR, and FOS at EC_50_ concentrations for 12 hours at 30 hpi. The positive control was PYR and the inactive negative control was FOS. The vehicle controls were at 0.5% (GA, XL, and PYR: DMSO and FOS: PBS). Relative DNA (SYBR assay) was normalized to corresponding vehicle controls. Data shown as means ± SEM, *n*=3 biological replicates; not significant (ns) >0.05, **<0.01, ****<0.0001; Welch’s t-test. (C-D) *P. falciparum*-infected RBCs were incubated with 10 µM EdA and drugs at EC_50_ concentrations for 12 hours at 30 hpi, fixed at 42 hpi, and stained. (C) Representative confocal microscopy images of parasites stained with Alexa Fluor Azide 488 (green) to label newly synthesized DNA and Hoechst (blue) to label nuclei. Yellow arrows highlight infected cells. Scale bars are 5 µm. (D) The integrated density of EdA signal demonstrates decreased EdA incorporation with GA, XL, and PYR treatments. Median indicated with red bar, *n*=3 biological replicates analyzing >300 parasites for each condition; not significant (ns) >0.05, **<0.01, ***<0.001; Mann-Whitney U test.

To next quantify changes in total DNA content following PfHsp90 inhibition, an adapted SYBR Green assay was employed (**Figure 4B**). Briefly, *P. falciparum*-infected RBCs were incubated with compounds at the times and concentrations indicated above, lysed with buffer containing SYBR Green I nuclear stain, and its signal was quantified and normalized to a vehicle control. Pyrimethamine (PYR), a dihydrofolate reductase (DHFR) inhibitor that halts DNA replication by depleting the nucleoside pool,^66,67^ was used as the positive control and FOS was used as the inactive negative control. Of note, FOS has been shown to impact parasite viability a cycle after its administration, thus *P. falciparum* inhibition is not expected by this compound under the experimental conditions.^68,69^ Consistent with our prior viability assessment (**Figures 1D-E**), 8- hour drug incubation at 30 hpi yielded no significant changes in relative DNA content following GA, XL and PYR treatments at both concentrations (**Figure S3B**). However, following the 12-hour incubation that captures DNA replication during schizogony, DNA signal of GA- and XL-treated parasites was significantly reduced, similar to the PYR control. At EC_50_ concentrations the signal was 32–38% reduced (**Figure 4B**), with a more pronounced reduction at 10xEC_50_ concentrations (**Figure S3C**). In contrast, the two vehicle controls (DMSO and PBS) and the inactive negative control (FOS) showed no significant change in DNA signal at either concentration after 12 hours. These observations suggest that the PfHsp90 inhibitors GA and XL likely disrupt parasite genome amplification during schizogony.

In 2022 Botnar *et al*. reported successful incorporation of alkyne-modified nucleosides into the newly synthesized DNA of *P. falciparum* with minimal cytotoxicity, enabling fluorescent detection via copper-catalyzed click chemistry (**Figure S4A**).^70^ Since SYBR Green staining does not differentiate between pre-existing and nascent DNA, we sought to apply this chemical tool to probe the impact of PfHsp90 inhibition on DNA synthesis. In this experiment, *P. falciparum*-infected RBCs were incubated with 10 µM 7-deaza-7-ethynyl-2’-deoxyadenosine (EdA) in the presence of compounds at the EC_50_ and 10xEC_50_ concentrations at 30 hpi for 12 hours. The cells were then fixed at 42 hpi, stained with 12 µM Azide-fluor 488 for bioconjugation to incorporated EdA and Hoechst to label parasite nuclei, and then assessed by confocal microscopy. To first ensure the specificity of EdA incorporation and click labeling, staining without EdA or Azide-fluor 488 was conducted, where no signal was detected, confirming the specificity of the approach (**Figures S4B** and **S4C**). We then assessed staining of EC_50_ concentration treatments and observed a notable reduction in the EdA signal per parasite, with less distinctly outlined and more diffuse nuclei with GA-, XL-, and PYR-treatment (**Figure 4C**). This phenotype was more pronounced with 10xEC_50_ treatment of these compounds (**Figure S5A**). In contrast, treatments with an inactive negative control (FOS), which was not expected to exhibit activity with this treatment duration, and vehicle controls (DMSO and PBS) retained round, well-defined nuclei morphology with discrete EdA signal. To quantitate compound-dependent changes in the degree of EdA incorporation, the integrated EdA density was measured for each condition, yielding a significant decrease after GA-, XL-, and PYR-treatment at both concentrations, but no change with FOS (**Figures 4D** and **S5B**).

EdA incorporation in *P. falciparum* was next tested with a small panel of fast-acting antimalarials with distinct mechanisms of action to explore the specificity of the phenotype. Experiments were completed as described above but with EC_50_ concentrations of chloroquine (CQ), which hinders heme detoxification,^71^ dihydroartemisinin (DHA), which generates reactive radicals among other outcomes,^72,73^ and cipargamin (CIP) that targets Na^+^ efflux.^74^ While XL reduced EdA incorporation, a decrease was observed with all three antimalarials (**Figures S6A** and **S6B**). Thus, diverse mechanisms of action can ultimately lead to inhibition of DNA replication, which is detectable by the EdA incorporation assay. In regard to XL and GA, the observed physical association of PfHsp90 with DNA replication components by co-IP supports a functional link between these pathways.

### Hsp90 inhibition disrupts nascent DNA synthesis during the *P. berghei* liver stage

To probe the *Plasmodium* stage specificity of the Hsp90-DNA synthesis connection, we employed a widely used rodent *P. berghei* liver stage model. For this, *P. berghei*-infected HepG2 hepatocytes were treated with EdA and inhibitors for 6 hours during 24–30 hours post-infection, capturing the onset of schizogony. EdA incorporation into actively replicating *P. berghei* parasites was validated prior to the experiment, where no significant EdA signal was observed in the absence of EdA (**Figures S7A** and **S7B**). Since previous studies reported reduced Hsp90 inhibitor potency when administered at 24 hours post-infection in *P. berghei*, we assessed changes in EdA incorporation following treatments with GA and XL at 5x and 50x their EC_50_ concentrations.^12,13^ We observed distinct nuclear morphology of EdA in the DMSO vehicle control, with a prominent reduction in EdA signal following GA, XL, and PYR treatments at both concentrations (**Figures 5A** and **S8A**). Quantification of the mean EdA intensity per parasite area revealed a >2-fold reduction following 5xEC_50_ and 50xEC_50_ GA and XL treatment, suggesting reduced DNA replication (**Figures 5B** and **S8B**). We additionally noted a significant decrease with the PYR positive control, although this reduction was more modest when compared to the tested Hsp90 inhibitors. Concurrently, host toxicity was visually assessed following the 6-hour incubation with compounds, where no impact on nuclei counts was observed. These observations align with the previously reported absence of HuH7 cytotoxicity after a 48-hour incubation with GA and XL up to 10 µM.^12,13^ Other liver stage inhibitors were next assessed for their impact on EdA incorporation, including cyclosporin A (CSA), which is inactive during the treatment window, and cycloheximide (CX), an established late liver stage translation inhibitor (**Figures S9A** and **S9B**).^75^ Following 5x and 50xEC_50_ CSA treatment, no changes in EdA incorporation were observed, as expected. Interestingly, CX treatment at the 5xEC_50_ concentration did not significantly impact the EdA signal, but it was significantly reduced at the 50xEC_50_ concentration. Thus, consistent with the blood stage observations, EdA incorporation in liver stage parasites can be impacted by molecules with different mechanisms of action.

**Fig 5.**
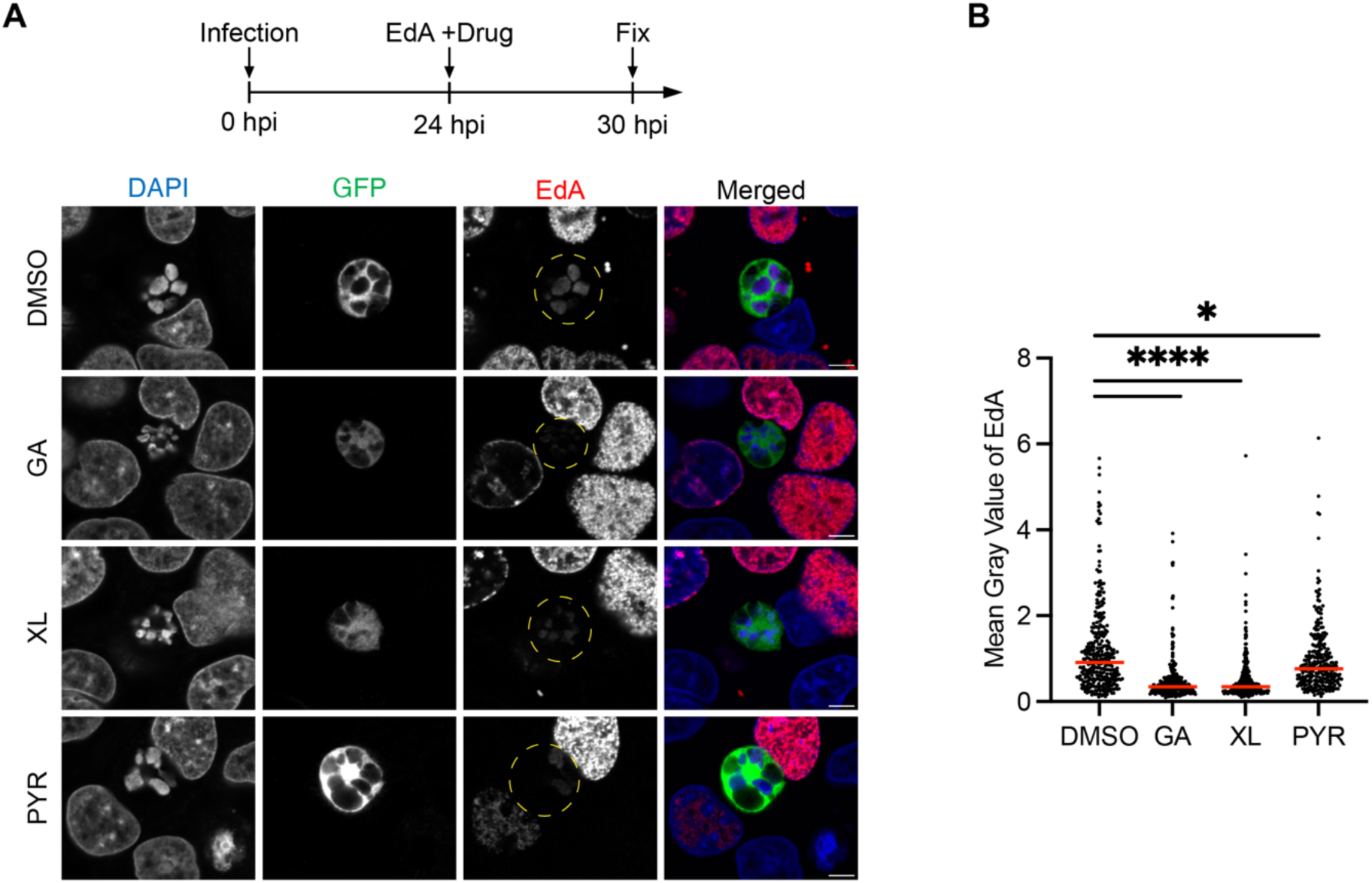
Inhibition of Hsp90 impairs DNA replication of *P. berghei* liver stage. (A-B) HepG2 cells were infected with GFP-expressing *P. berghei*, treated with 5 µM EdA and drugs at 5xEC_50_ concentrations for 6 hours at 24 hours post-infection (hpi), fixed at 30 hpi and stained. (A) Representative confocal fluorescence microscopy images of parasites stained with anti-GFP (green) to label parasites, AZDye594-Azide (red) to label newly synthesized DNA, and DAPI (blue) to label nuclei. Yellow dashed circles highlight infected cells. Scale bars are 5 µM. (B) The mean gray value of EdA demonstrates decreased EdA incorporation following GA, XL, and PYR treatments. Median indicated with red bar, *n*=3 biological replicates analyzing >250 parasites for each condition; *<0.05, ****<0.0001; Mann-Whitney U test.

### Targeting PfHsp90 sensitizes *P. falciparum* to DNA replication inhibitors

Drug synergy is critical for combating parasite drug resistance, which has eroded the efficacy of all current antimalarials. Building on the observed connection between PfHsp90 activity and DNA replication, we questioned whether PfHsp90 inhibition could sensitize parasites to DNA replication inhibitors. Synergy was evaluated based on parasite viability and analyzed using SynergyFinder 3.0,^76,77^ which has been previously used to assess synergy for anti-*Plasmodium* and anti-cancer agents.^13,78–83^ The program assigns a synergy score for each drug pair and categorizes interactions as antagonistic (score<-10), additive (-10 to 10), or synergistic (>10).

Using this workflow, we first assessed synergies via the zero interaction potency (ZIP) reference model which compares the potency shifts between combination treatments and their monotherapies across dose-response curves. The synergistic combination of artesunate (AS) and mefloquine (MQ), the frontline ACT therapy against malaria, was assessed as a positive control, where many synergistic interactions with scores ranging from 10 to 60 were observed (**Figure S10B**). We also assessed synergy with a more classical Bliss model which assumes independent drug action and does not account for dose-response relationships.^84^ Synergy was also identified for AS and MQ with this model (**Figures S11A** and **S12A**).

We next assessed the combination of Hsp90 and DNA replication inhibition by employing three compounds that target distinct components of the DNA replication machinery: aphidicolin (APH) which inhibits DNA polymerase α, δ, and ε;^85^ etoposide (ETOP) which blocks topoisomerase II (TOP2);^86^ and PYR which reduces nucleoside levels through DHFR inhibition (**Figures 6A-C**).^66^ The EC_50_ concentrations of the compounds against *P. falciparum*-infected RBCs were determined using the aforementioned SYBR Green assay. APH, ETOP and PYR inhibited *P. falciparum* with EC_50_s of 0.98 ± 0.016 µM, 4.96 ± 0.51 µM and 0.034 ± 0.0019 µM, respectively (**Figure S10A**). The EC_50_ values of APH and PYR are consistent with previous reports,^67,87^ whereas ETOP is an established PfTOP2 inhibitor for which we report the blood stage EC_50_ in this study.^88^ Under the ZIP model, compound co-treatments produced localized synergy at the EC_50_ concentration of XL and sub-EC_50_ concentration of APH (**Figures 6D**), while co-treatment of XL with ETOP led to several synergistic regions at sub-EC_50_ concentrations of both compounds (**Figures 6E**). For these combinations, the scores ranged up to 12, indicating moderate synergy. XL synergy with APH and ETOP was also detected under the Bliss model, suggesting the results are robust and likely independent of the synergy assessment method (**Figures S11B-C, S12B-C**). In contrast, no synergistic interactions were observed with XL and PYR co-treatments with either the ZIP or Bliss model (**Figures 6F**, **S11D,** and **S12D**). The synergistic interaction between XL and two distinct DNA synthesis inhibitors provides further support of a functional link between PfHsp90 and DNA replication.

**Fig 6.**
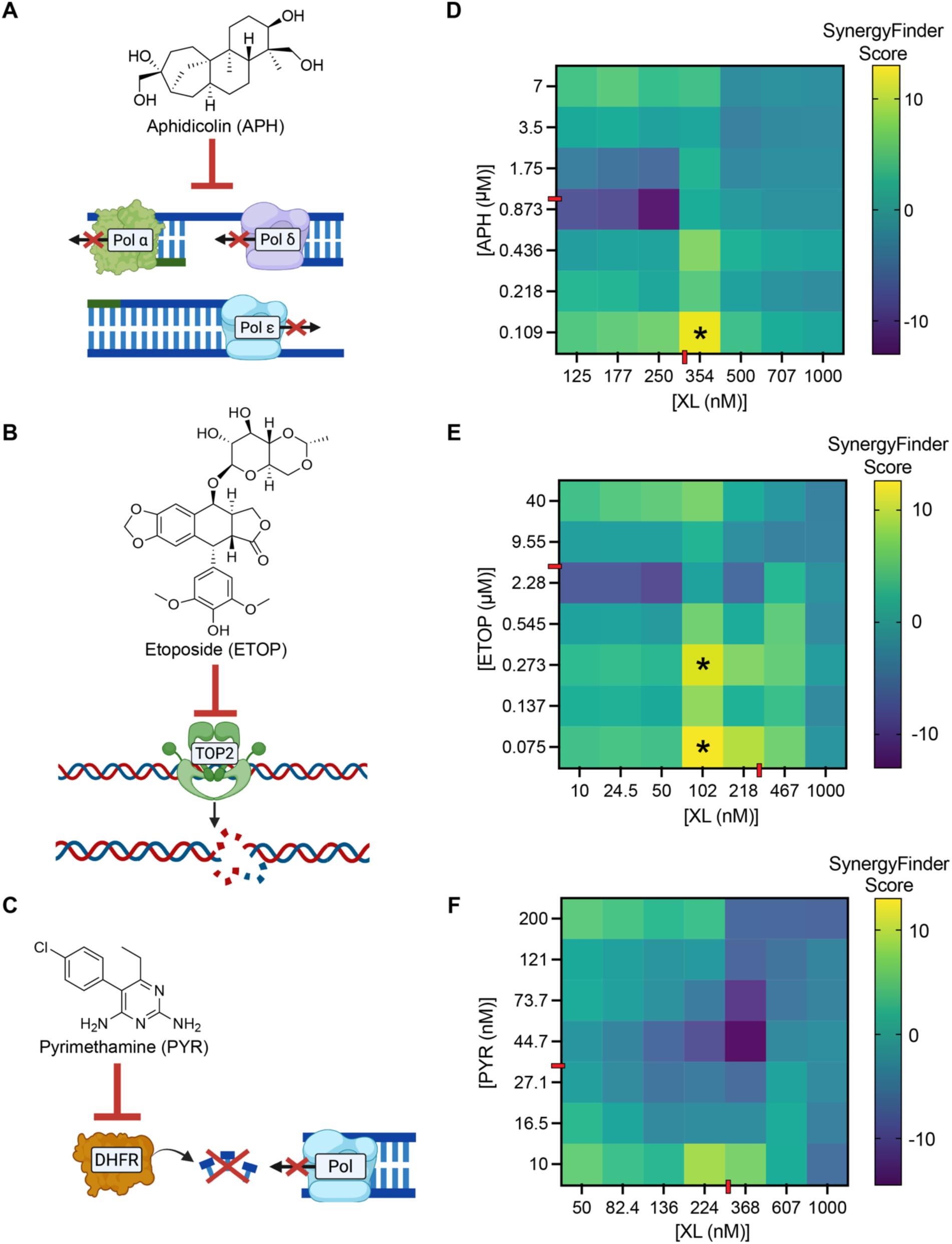
PfHsp90 inhibition sensitizes *P. falciparum* to DNA replication inhibitors. (A-C) Modes of action of the DNA replication inhibitors (A) aphidicolin (APH), (B) etoposide (ETOP) and (C) pyrimethamine (PYR). (D-F) Synergy heatmaps depicting combined effects of XL with APH (D), ETOP (E), and PYR (F). Synergistic interactions (SynergyFinder score>10) are marked with an asterisk (*). The red dashes indicate EC_50_ concentrations for each compound. Data shown as means, *n*=3 biological replicates.

We next sought to examine if XL synergizes with mechanistically diverse compounds. Specifically, we tested combinations of XL with FK506, a FKBP35 inhibitor previously shown to synergize with Hsp90 inhibitors in *Plasmodium*,^89^ as well as CQ, DHA, and CX. As expected, we observed a synergistic interaction at sub-EC_50_ concentrations of FK506 and XL under both the ZIP and Bliss models (**Figures S10C**, **S11E**, and **S12E**). We further noted a synergistic interaction of XL with DHA, while no synergy was observed with CQ and CX under either model (**Figures S10D-F, S11F-H** and **S12F-H**).

## DISCUSSION

In the deadliest malaria parasite, *P. falciparum*, Hsp90 is a key regulator of proteostasis which has been recognized as a putative drug target for over two decades due to its critical role in parasite survival and pathogenicity.^11,31^ Despite this, our understanding of PfHsp90-dependent proteins and how its chemical perturbation reshapes parasite biology has been limited. Aiming to address this, we leveraged global profiling of PfHsp90 inhibition to uncover parasite pathways supported by the chaperone and investigate possible interactors. We identified 131 putative high confidence PfHsp90-dependent proteins, where a co-IP study further supported a physical interaction with 52 of these proteins. Specifically, bioinformatic analyses highlighted a link between Hsp90 and DNA replication. In support of this proposal, we found that Hsp90 disruption not only leads to a reduction in the total parasite DNA but also impairs nucleoside incorporation during the blood and liver stages. Beyond this, our rich datasets provide hundreds of compelling starting points that could be the focus of future work. Significantly, we observed that PfHsp90 disruption sensitizes *P. falciparum* to DNA replication inhibitors, revealing a therapeutic value for the functional link between the two pathways.

Studies examining the impact of Hsp90 perturbation on the proteome have been instrumental in revealing the complex roles of the chaperone by providing meaningful physiological insights extending beyond an immediate client network.^90^ Such studies have identified previously unrecognized Hsp90-dependent kinases in cancer cell lines, uncovered resistance-associated biomarkers in prostate tumors, and demonstrated the regulatory role of Hsp90 in nascent protein synthesis and expression of virulence factors in *Leishmania* parasites.^35,36,91^ This approach may also detect Hsp90 clients, although the methodology does not directly measure an interaction. To the best of our knowledge PfRad51 and PfSir2A are the only two validated PfHsp90 clients,^29,30^ and these were not among our Hsp90-dependent proteins. To directly inform proteins that may physically interact with Hsp90, we complemented our analyses with co-IP profiling and mining of well-established yeast and human interactome datasets. Notably, the Hsp90 client PfRad51 was detected by our co-IP.

In 2013 Shahinas and colleagues^28^ performed a PfHsp90 co-IP coupled with LC-MS/MS, identifying 186 putative PfHsp90 interactors. However, direct comparison of our PfHsp90-dependent proteins with that study was precluded, as a significant portion of their pulled down proteins lacked annotation, likely reflecting limitations in the MS technology and database annotations of the time. Our co-IP identified 52 PfHsp90-dependent proteins that may interact with the chaperone. Of note, co-IP does not differentiate between direct and indirect interactors of PfHsp90, as proteins may be pulled down as a part of larger complexes. To predict direct physical interactions with PfHsp90, we attempted to employ the AlphaFold3 server;^92^ however, even for the two PfHsp90 clients the method yielded low interface predicted template modeling (ipTM) scores, likely due to the intrinsically disordered nature of PfHsp90’s charged linker region (**Figure S13**). Further complicating predictions, there is no general conserved motif among Hsp90 clients.^93^ Past investigations have established that yeast Hsp90 associates with intrinsically disordered regions (IDRs) to regulate client function.^18,94^ In an effort to evaluate client-like features within our dataset, we examined enrichment of IDRs and disordered binding regions (DBRs), subsets of IDRs capable of undergoing disorder-to-order transition upon partner interaction.^95^ Our efforts yielded no IDR or DBR enrichment when compared to randomly pooled parasite proteins, a result that was not surprising given that one-third of the *P. falciparum* proteome is predicted to be disordered (**Figure S14, Table S3D**).^96^ This underscores the importance of expanding the number of validated *bona fide* clients by directly testing if our identified proteins bind to PfHsp90 and are refolded. Additionally, because our protein identification relied on compromised stability and thus degradation upon PfHsp90 inhibition, transcriptomic profiling after GA or XL treatment would provide a complementary approach to distinguish direct PfHsp90-sensitive partners from secondary responses through comparison of protein- and transcript-level changes.

PfHsp90-dependent processes have been largely understudied until a recent TPP investigation that uncovered a link between Hsp90 and the proteasome.^13^ This important starting point shed light on putative PfHsp90-sensitive proteins, but weaker transient molecular partners may have been missed since it was conducted in parasite lysate and employed a mass spectrometry method inherently biased toward higher abundance proteins (DDA). Our approach overcomes these limitations by performing compound treatment in live parasites, capturing changes resulting from PfHsp90 inhibition in the native environment. We further leveraged a DIA proteomic approach to ensure coverage of low-abundance proteins, achieving one of the highest *P. falciparum* proteome coverages reported to date.^51,52^ Among our PfHsp90-dependent proteins, 4 (Q8I3J5, Q8IL94, Q8ILR7 and Q9U0J0) were previously identified in the TPP study that leveraged the resorcinol-based inhibitor, BX-2819. Beyond this direct overlap, we observe an additional overlap of 7 proteins when considering components of hetero-oligomeric complexes, underscoring the ability of our approach to identify PfHsp90-sensitive proteins. This 22% overlap likely reflects differences in experimental design and methodology, specifically utilizing live parasites over lysate, varying drug incubation times, MS approaches and confidence assignment cut-offs. Importantly, proteins involved in DNA replication were represented in all distinct proteomic analyses (PfHsp90-dependent, co-IP, and TPP).

Unlike most organisms, *Plasmodium* parasites do not replicate through traditional binary fission and instead undergo schizogony, where multiple rounds of replication produce 16 – 32 daughter cells during the blood stage.^97^ The success of *P. falciparum* pathogenicity is linked to its high multiplication rates.^98^ Coupled with the absence of several human DNA replication proteins and canonical cell cycle checkpoints, *P. falciparum* DNA replication represents a compelling target for antimalarial intervention.^99–101^ In this study we found that parasite treatment with PfHsp90 inhibitors during the trophozoite-to-schizont transition results in DNA reduction. While decreased DNA levels could reflect a delay or stall in stage development, our finding that GA- and XL-treated parasites progress to schizonts supports partial replication inhibition rather than a complete arrest. The lack of rise in rings, even by 42 hpi, also suggests that PfHsp90 inhibition likely disrupts schizogony itself, thereby preventing egress and reinvasion, rather than simply delaying the stage progression. Previous cancer studies have shown that transient Hsp90 inhibition decreases active DNA synthesis, reflected by reduced 5-ethynyl-2′-deoxyuridine (EdU) incorporation.^61^ We completed the complementary study using an adenosine analog EdA, previously shown to natively incorporate into *Plasmodium* nascent DNA during the blood and liver stages,^70^ and applied it for the first time to study drug-induced perturbations on parasite DNA replication. We observed a significant decrease in EdA incorporation in GA- and XL-treated parasites during both stages. Importantly, our assessment of mechanistically diverse compounds demonstrates that reduced nucleoside incorporation can result from either direct DNA replication inhibition or parasite stress that indirectly impairs replication. However, the fact that numerous components of polymerases, primases, MCM2-7 complex, and other replication proteins were significantly reduced following PfHsp90 inhibition and enriched by co-IP, supports the functional link between replication machinery and PfHsp90. We speculate that PfHsp90 associates with the replisome and likely supports the stability and activity of its constituent proteins. This further aligns with the well-established role of Hsp90 chaperones in regulating DNA replication across other diverse organisms.^61,102,103^

We investigated the major cluster from our analysis – the 68 proteins that mapped to the DNA replication pathway in the PPI network. The second largest cluster was amino sugar and nucleotide sugar metabolism, where three proteins (Q8IDQ3, Q8II63 and Q8ILP1) are enzymes involved in glycosylphosphatidylinositol (GPI) anchor biosynthesis, which promotes *P. falciparum* pathogenicity through induction of host inflammatory responses and anchorage of merozoite surface proteins.^104–106^ Intriguingly, previous studies established that anti-GPI antibodies confer protection against malaria and that targeting GPI anchors abrogates *P. falciparum* growth.^107,108^ Building on this, our findings suggest that PfHsp90 might play a role in supporting the stability and function of multiple GPI anchor biosynthetic enzymes, providing a promising molecular link that has yet to be explored.

Combination therapies have been instrumental in malaria control efforts with ACTs serving as the frontline treatment for decades. Pairing compounds with distinct mechanisms of action has proven advantages, including lowered individual drug doses, reduced toxicity, increased parasite clearance and delayed resistance.^109,110^ However, the recent decline in *P. falciparum* susceptibility to ACTs highlights the pressing need to identify novel drug combinations.^111^ Building on our findings, we hypothesized that co-treatment of DNA replication inhibitors with the clinically favorable XL could potentiate antimalarial activity. To assess synergy we utilized SynergyFinder, which surveys full dose-response matrices across multiple reference models, while the more traditional fractional inhibitory concentration (FIC) approach assays fixed single-point concentration doses.^112^ Among the tested DNA replication inhibitors, APH was particularly interesting since it targets α, δ, and ε DNA polymerases,^85^ all of which were among our PfHsp90-dependent and pulled down proteins. We were also interested in the FDA-approved drug ETOP given its previously established synergy in cancer cell lines with 17-AAG.^113,114^ Our efforts demonstrated synergistic interactions for APH and ETOP co-treatments with XL, further supporting a functional link between PfHsp90 and DNA replication. Moreover, this finding exposes a vulnerability in the parasite that may be leveraged in future drug development efforts. For these efforts, DNA replication inhibitors must be designed to avoid harming host machinery, a consideration that was not needed when targeting *Plasmodium* within anucleated RBCs. Further, improving selectivity for PfHsp90 over HsHsp90 remains a pressing challenge for the development of viable therapeutic agents and improved chemical probes.

Overall, our in-cell PfHsp90 inhibition and co-IP analyses highlight the importance of Hsp90 for *Plasmodium* DNA replication and point to compelling biological processes to explore in future studies. These candidates include proteins conserved as Hsp90 interactors across species, as well as proteins unique to *Plasmodium*, where the latter represent promising targets for future therapeutic development and the discovery of novel Hsp90 functions. Our datasets also provide a critical starting point for future studies aimed at identifying PfHsp90 clients, whose functional dependency on PfHsp90 may be exploited in future antimalarial combination therapies. Ultimately, our efforts underscore the value of continued investigation of PfHsp90 and its functional network, offering a promising avenue for future antimalarial drug discovery.

### Limitations of the study

One limitation of our study is the susceptibility of proteomic analysis to false positives and negatives, coupled with the potential for small-molecule off-targets. Future studies exploring GA and XL off-targets will be critical to deconvolute their full target spectrum. To this point, the known targeting of HsHsp90 highlights the need to develop PfHsp90 inhibitors with improved species-selectivity and biophysical properties to support cellular studies. Such molecules could be used to refine our PfHsp90-dependent proteome and probe PfHsp90-specific phenotypes. As another limitation, Hsp90-dependent processes that function at distinct stages or upon heat stress could have been missed due to our experimental workflow. Further, our synergy assays do not address whether co-treatment with DNA replication and Hsp90 inhibitors can selectively target *P. falciparum* over host cells. Building on this proof-of-principle study, the next step is to develop inhibitors that selectively target the parasite DNA replication machinery to maximize therapeutic selectivity and minimize host toxicity. Lastly, while our co-IP analysis identifies candidate PfHsp90 interactors, the co-IP itself does not distinguish between a direct or indirect interaction. Thus, future efforts employing biochemical orthogonal assays will be critical in validating interactors or clients.

## STAR★Methods

### RESOURCE AVAILABILITY

#### Lead contact

Further information and requests for resources and reagents should be directed to and will be fulfilled by the lead contact, Emily R. Derbyshire.

#### Materials availability

This study did not generate new unique reagents.

#### Data and code availability

The data have been deposited to the ProteomeXchange Consortium via the PRIDE partner repository with the dataset identifier PXD079493 for inhibition-based study and to MassIVE repository with the dataset identifier PXD080188 (MassIVE MSV000102274) for co-IP.

### MATERIALS AND METHODS

#### Experimental model and study participant details

##### Parasite Lines

*P. falciparum* 3D7 (BEI Resources, NIAID, NIH, MRA-102) was maintained in 10.44 g/L RPMI 1640 (Gibco), 25 mM HEPES (Gibco), pH 7.2, 0.37 mM hypoxanthine (Sigma), 24 mM sodium bicarbonate (Sigma), 0.5% (wt/vol) AlbuMAX II (Gibco) and 25 µg/mL gentamicin (Sigma) at 37 °C with 5% CO_2_. Whole blood (Gulf Coast Regional Blood Center, Houston, TX) was used to supplement parasites with fresh erythrocytes. Every 48 hours (hrs), parasites were synchronized at the ring stage using 5% (wt/vol) D-Sorbitol (Sigma).

HepG2 cells (ATCC HB-8065, Duke Life Science Facility) were maintained in Dulbecco’s Modified Eagle Medium (DMEM, Gibco) supplemented with 10% (vol/vol) fetal bovine serum (Millipore Sigma) and 1% (vol/vol) antibiotic-antimycotic (Millipore Sigma) in a standard tissue culture incubator at 37 °C with 5% CO_2_. *Anopheles stephensi* mosquitoes infected with *P. berghei* ANKA stably expressing GFP (GFP-expressing *P. berghei*) were obtained from the SporoCore University of Georgia, Athens. *P. berghei* sporozoites were isolated from mosquito salivary glands before infections.

##### Bacterial Strains

*Escherichia coli* NEB5α (NEB) and BL21 (DE3) (NED) were utilized for plasmid and protein expression, respectively. Bacteria were propagated via Difco^TM^ Luria-Bertain (Miller) agar plates and cultured in Difco^TM^ Luria-Bertain (Miller) broth supplemented with Ampicillin (Sigma) antibiotic.

### Method details

#### Protein expression and purification

For protein expression, *E. coli* BL21 (DE3) cells (New England Biolabs) were transformed with pET-21 PfHsp90-His_6_ and pET-21 HsHsp90-His_6_ and grown in 1 L of Luria-Bertani broth containing 100 µg/mL ampicillin (Sigma) at 37 °C with shaking at 250 rpm. At an optical density of 0.6 – 0.8 at 600 nm, the temperature was reduced to 18 °C and the cultures were induced with 100 µM isopropyl-ß-D-1-thiogalactopyranoside (IPTG). After 18 hrs at 250 rpm, cell pellets were collected by centrifugation at 4,300 *g* and stored in -80 °C. For cell lysis, pellets were resuspended in buffer A (50 mM KH_2_PO_4_, pH 8.0, 200 mM NaCl, 1 mM benzamidine, 5% glycerol (vol/vol), 5 mM β-Mercaptoethanol, 10 mM Imidazole supplemented with 1 cOmplete, Mini, EDTA-free Protease Inhibitor Cocktail tablet (Roche)). The pellets were lysed by 6 rounds of sonication (Fisher Scientific), centrifuged, and the supernatant was incubated with Ni-NTA agarose resin (Qiagen) overnight at 4 °C. Following stringent washes with buffer A the protein was eluted in a linear imidazole gradient (25-150 mM). Fractions containing the protein of interest were exchanged into buffer (25 mM TEA, pH 8.0, 25 mM NaCl, 1 mM DTT, 5% glycerol (vol/vol)) via 30K MWCO centrifugal device (Pall corporation) and stored in -80 °C. The protein concentration was determined via Pierce Coomassie Plus Bradford Assay Reagent (ThermoScientific).

#### *P. falciparum* blood stage culture

The wild-type *P. falciparum* 3D7 parasites (MRA-102, BEI resources) were continuously cultured *in vitro* in a complete medium (10.44 g/L RPMI 1640 (Gibco), 25 mM HEPES (Gibco), pH 7.2, 0.37 mM hypoxanthine (Sigma), 24 mM sodium bicarbonate (Sigma), 0.5% (wt/vol) AlbuMAX II (Gibco) and 25 µg/mL gentamicin (Sigma)). Cultures were maintained at 37 °C with 5% CO_2_. Every 48 hrs the parasites were synchronized with 5% (wt/vol) D-Sorbitol (Sigma) and supplemented with freshly washed human erythrocytes (Gulf Coast Regional Blood Center, Houston, TX).

#### *P. falciparum* blood stage inhibition assays

*Total DNA content. P. falciparum* parasites were synchronized as described above and adjusted to 10% parasitemia at 1% hematocrit. Parasites at the mid-trophozoite stage (30 hpi) were added to a UV-sterilized black 96-well non-treated plate (Corning). The compounds were administered at their literature EC_50_ and 10xEC_50_ concentrations. GA (ApexBio) and XL888 (ApexBio) dissolved in DMSO (Sigma) were each administered at 0.3 and 3 µM. Pyrimethamine (Cayman Chemical) dissolved in DMSO was administered at 0.05 and 0.5 µM. Drugs were added to parasite cultures in each well to a final volume of 200 µL, with a final DMSO concentration of 0.5%, which was used as the negative control. Fosmidomycin (Cayman Chemical) was administered at its reported EC_50_ concentration of 1.17 µM and 10xEC_50_ concentration of 11.7 µM in phosphate-buffered saline (PBS) (ThermoFisher Scientific). For fosmidomycin assessment, 0.5% PBS was the negative control. Following compound addition, parasites were incubated at 37 °C in 5% CO_2_ for either 8 or 12 hrs. Parasite total DNA content was then assessed by addition of 40 µL of lysis buffer (20 mM Tris-HCl pH 7.5 (Fisher Chemical), 5 mM EDTA dipotassium salt dihydrate (Fisher Chemical), 0.16% (wt/vol) saponin (Sigma), and 1.6% (vol/vol) Triton X-100 (Fisher Chemical)) with 10x SYBR Green I (Invitrogen) to each well. After an overnight incubation at room temperature (RT) in the dark, relative parasite DNA was quantified by measuring the fluorescent signal at an emission of 535 nm and excitation of 485 nm using an EnVision 2015 multimode plate reader (PerkinElmer). Relative DNA content was calculated by normalizing the fluorescence signal to the negative controls. The experiment was performed in six technical replicates across three biological replicates.

*Stage progression*. *P. falciparum* parasites were synchronized as described above and adjusted to 10% parasitemia at 1% hematocrit. At the mid-trophozoite stage (30 hpi) parasites were added to a sterile 12-well non-treated plate (Corning). GA and XL were added at their EC_50_ and 10xEC_50_ concentrations as described above. Drugs were added to parasite cultures in each well to a final volume of 1.5 mL, with a final DMSO concentration of 0.5%, which was used as a negative control. E64d (Cayman Chemical) was administered at its reported EC_50_ of 0.6 and 10xEC_50_ concentration of 6 µM. Following compound addition, parasites were incubated at 37 °C in 5% CO_2_ for either 8 or 12 hrs. Following incubation, blood smears were obtained and stained (Hemacolor kit, Sigma). The number of rings, trophozoites, and schizonts were counted for each smear for a total of >290 parasites per treatment at each timepoint, across 3 biological replicates. *Synergy. P. falciparum* parasites were synchronized as described above and adjusted to 2% parasitemia at 1% hematocrit. Parasites at the ring stage (10-14 hpi) were added to a UV-sterilized black 96-well non-treated plate (Corning). XL, aphidicolin (Cayman Chemical), etoposide (Cayman Chemical) and pyrimethamine were administered via a HP D300 Digital Dispenser. Quinacrine (Sigma) at 0.5 µM was used as the positive control while 0.5% DMSO was the negative control. Plates were incubated at 37 °C 5% CO_2_ for 72 hrs. Following incubation, parasite viability was evaluated as described above. Data was normalized to the negative (0,0 point; 0.5% DMSO) and positive (0.5 µM quinacrine) controls. EC_50_ values were obtained by fitting data to a standard dose response equation (GraphPad Prism). Synergy was evaluated via SynergyFinder 3.0 zero interaction potency (ZIP) and Bliss reference models with baseline correction. The experiments were performed in three technical replicates for APH, ETOP, PYR, FK506, CQ, and DHA, and two technical replicates for AS and MQ, across three biological replicates for all drug combinations. The Z-factor ranged from 0.5 to 0.9.

#### Inhibitor treatment and parasite isolation for proteomic analysis

Mid-trophozoite stage (30 hpi) *P. falciparum* parasites were grown at 7% parasitemia and 1% hematocrit. For each replicate a culture was split into 80 mL for drug treatments. The cultures were subsequently incubated with 0.3 µM GA, 0.3 µM XL, or 0.5% DMSO for 8 hrs at 37 °C and 5% CO_2_. Prior to and after the drug treatment until the trophozoite–mid trophozoite stage (38 hpi), the parasitemia was quantified for each treated culture (Hemacolor staining kit). For this analysis, a total of 900 – 1500 RBCs were evaluated per replicate before and after each treatment. After 8 hrs, parasites were subsequently pelleted by centrifugation and released from host RBCs via the addition of 0.03% (wt/vol) saponin in PBS. Released parasites were pelleted by centrifugation and washed 4 times with cold PBS to ensure the removal of host RBCs and lysis debris. Pellets were stored at -80 °C until further processing for downstream proteomic analysis. Inhibitor treatment was performed across three biological replicates for all treatment conditions, with the first DMSO replicate performed in technical duplicates.

#### Co-immunoprecipitation

*P. falciparum* parasites were grown to 10% parasitemia and 1% hematocrit. At the late-trophozoite stage (38 hpi), parasites were isolated from RBCs as described above and processed immediately. The co-IP was performed with Pierce™ Crosslink Magnetic IP/Co-IP Kit (88805) for anti-PfHsp90 antibody (Sigma) and an anti-IgG antibody as a control (ThermoFisher Scientific). Parasite pellets were lysed in lysis buffer (IP Lysis/wash buffer from the kit supplemented at 1:7 with cOmplete, Mini, EDTA-free Protease Inhibitor Cocktail tablet) for 30 min. The lysates were subsequently clarified by centrifugation and protein concentrations were quantified by Bradford Assay. Protein A/G magnetic beads (Pierce) at 25 µL of bead slurry were pre-washed with 1x Modified Coupling Buffer and conjugated to the antibodies for 15 min. The beads were subsequently washed 3 times with 1x Modified Coupling Buffer and 2 times with IP Lysis/Wash buffer. Following this, 500 µg of lysate was incubated with antibody-conjugated beads for 1 hr at 4 °C. The beads were washed 3 times with cold IP Lysis/Wash buffer and ultrapure water. The proteins were eluted with boiling for 5 min in elution buffer (1.5% SDS and 10 mM DTT in PBS) and flash-frozen.

#### Sample preparation for LC-MS/MS

##### Inhibition-based proteomics

Parasite pellets were reconstituted in 4 volumes of PBS supplemented with protease inhibitor cocktail (1 mM AEBSF, 20 μM leupeptin, 10 μM pepstatin A, 500 μM bestatin, and 15 μM E-64) in 1:100 cocktail to buffer ratio. Pellets were lysed by sonication, centrifuged, and the supernatant was analyzed via bottom-up proteomic analysis. The protein concentration was assessed via Bradford Assay. Samples were analyzed across three to four biological replicates. Lysates from compound and DMSO-treated parasites were distributed into 10 kDa MWCO filters (Millipore), with each filter containing ∼80 μg of protein. The samples were washed 3 times with 300 µL 8 M urea (Sigma) in 0.1 M Tris-HCl (OmniPur) (pH 8.0), reduced with 5 mM TCEP (ChemCruz) at RT for 1 hr, alkylated with 20 mM S-methylmethanethiosulfonate (MMTS) (Sigma) for 15 min and then washed with 0.1 M triethylammonium bicarbonate (TEAB) (Supelco) buffer (pH 8.5) for 3 times. Proteins were digested with trypsin in 0.1 M TEAB buffer at 37°C for 16 hrs at a trypsin (ThermoFisher Scientific) to total protein ratio of 1:50. Resulting peptides were eluted with 50 µL of 0.5 M NaCl twice. 1/3 of the digested peptides were taken out for DIA after desalting using Silica C18 MacroSpin Columns (Higgins Analytical Inc) and vacuum-dried.

##### Co-Immunoprecipitation proteomics

Samples were spiked with 1 or 2 pmol bovine casein as an internal quality control standard. Next, they were brought to 5% SDS and reduced for 15 min at 80°C, alkylated with 20 mM iodoacetamide for 30 min at RT, then supplemented with a final concentration of 1.2% phosphoric acid and 695 µL of S-Trap (Protifi) binding buffer (90% MeOH/100 mM TEAB). Proteins were trapped on the S-Trap micro cartridge, digested using 1 µg sequencing grade trypsin (Promega) for 1 hr at 47°C, and eluted using 50 mM TEAB, followed by 0.2% formic acid (FA), and lastly using 50% ACN/0.2% FA. All samples were then lyophilized. Samples were resuspended in 200 µL of 0.1% FA. A study pool QC (SPQC) was created by combining equal volumes of each sample and run periodically throughout the study.

#### LC-MS/MS Analysis by Orbitrap Astral

##### Inhibition-based proteomics

LC-MS/MS used an Evosep One LC interfaced to a ThermoFisher Orbitrap Astral. The Evosep used a 30 sample-per-day method with a Bruker Pepsep 15 cm x 150 μm column (1.5 μm particle size) heated to 45 °C with a Butterfly portfolio heater and interfaced to a PepSep Sprayer and stainless steel (30 μm) emitter. MS/MS used a 240,000 resolution Orbitrap precursor scan every 0.6 s from m/z 380-980, an AGC target of 500% and 50 ms max fill time. DIA MS/MS scans were acquired in the Astral analyzer using fixed windows of 4 m/z from m/z 380-80 m/z, target AGC of 500% and max fill time of 6 ms. A normalized collision energy of 28% was used for all MS2 scans. Raw MS data was converted to *.htrms format using HTRMS converter and processed in Spectronaut 19. A spectral library was generated by direct-DIA searches of a Uniprot *H. sapiens* (downloaded on 9/13/2024; 20468 entries) and *P. falciparum* (downloaded on 07/18/2-24; 5479 entries) databases appended with common contaminants using FragPipe. Searches used fixed methylthiol modification on Cys and variable protein N-terminal acetylation and methionine oxidation, Trypsin/P specificity and up to 2 missed cleavages. A spectral library containing 131,219 precursors, 98,931 modified peptides and 6,204 protein groups was used for identification and quantification. Default settings were utilized except that the PTM workflow was selected.

##### Co-immunoprecipitation proteomics

Prior to LC-MS analysis, a fluorescent peptide level quantitation assay (Thermo) was performed on the SPQC sample and volumetric loading for all samples was adjusted such that a target 250 ng of the SPQC was loaded on each EvoTip (EvoSep). Additionally, 50 fmol of yeast ADH was loaded on each tip for QC purposes. Quantitative LC/MS/MS was performed using an EvoSep One UPLC coupled to a Thermo Orbitrap Astral high resolution accurate mass tandem mass spectrometer (Thermo). Briefly, each sample loaded EvoTip was eluted onto a 1.5 µm EvoSep 150 µm ID x 15 cm performance (EvoSep) column using the SPD30 gradient at 45 °C. Data collection on the Orbitrap Astral mass spectrometer was performed in a DIA mode of acquisition with a r=240000 (@ m/z 200) full MS scan from m/z 380-980 in the OT with a target AGC value of 5000000 ions. Fixed DIA windows of 4 m/z from m/z 380-980 DIA MS/MS scans were acquired in the Astral with a target AGC value of 50000 and max fill time of 6 ms. HCD collision energy setting of 28% was used for all MS2 scans. The total analysis cycle time for each sample injection was approximately 44 min.

#### Quantitative Data Analysis

##### Inhibition-based proteomics

Protein abundance data exported from Spectronaut 19 were processed and the protein intensities were globally scaled to adjust for variation in sample loading. The protein abundances were further normalized so that the mean abundance across all DMSO samples equaled 1. Statistical significance was assessed using one-way ANOVA followed by Bonferroni correction. For visualization, volcano plots were generated by plotting -log_10_ (*p*-values) against the average log_2_(fold change), with significance thresholds set at *p* <0.05 and fold change at >0.5 or <-0.5. Fold changes were calculated as GA/DMSO or XL/DMSO using the corresponding drug and DMSO pairs.

##### Co-immunoprecipitation proteomics

Following UPLC-MS/MS analyses, data were imported into Spectronaut v20 (Biognosys) and individual LCMS data files were aligned based on the accurate mass and retention time of detected precursor and fragment ions. Relative peptide abundance was measured based on MS2 fragment ions of selected ion chromatograms of the aligned features across all runs. The MS/MS data was searched against a reference proteome of *P. falciparum* (3D7 strain, downloaded in 2026), a SwissProt Human database (downloaded in 2026), a common contaminant/spiked protein database (bovine albumin, bovine casein, yeast ADH, etc.) and an equal number of reversed-sequence “decoys” for false discovery rate determination. A library free Direct DIA+ approach within Spectronaut was used to perform the database searches. Database search parameters included fixed modification on Cys (carbamidomethyl), variable modification on Met (oxidation) and N-term (acetylation). Full trypsin enzyme rules were used along with 10 ppm mass tolerances on precursor ions and 20 ppm on product ions. Spectral annotation was set at a maximum 1% peptide false discovery rate based on q-value calculations. The peptide homology was addressed using razor rules in which a peptide matched to multiple different proteins was exclusively assigned to the protein with more identified peptides. Protein homology was addressed by grouping proteins that had the same set of peptides to account for their identification. A master protein within a group was assigned based on % coverage. If less than half of the values are missing in a biological group, values are imputed with an intensity derived from a normal distribution of all values defined by measured values within the same intensity range (20 bins). If greater than half values are missing for a precursor in a group and a precursor intensity is greater than one standard deviation above the mean of all measured precursor intensities, then it was concluded that the peptide was misaligned, and its measured intensity is set to 0. All remaining missing values are imputed with the lowest 2% of all detected values. The IQ_MaxLFQ rollup strategy was deployed to convert from the precursor level to the protein level based on all remaining precursor values. Following database searching using a library free Spectronaut search, the data was annotated at a 1% peptide false discovery rate. There were 35,541 precursors identified in these datasets which corresponded to 3,102 total proteins. At least 2 unique peptides were required to be identified per protein and no missing values were required within any of the samples in one comparison group. Statistical significance was assessed with Benjamin Hochberg FDR corrected *p*-values. For visualization, volcano plots were generated by plotting -log_10_ (*p-*values) against the average log_2_(fold change), with significance thresholds set at *p*-value <0.05 and fold change at >5 and <-5.

#### Post-MS Data Analysis

##### Ortholog analysis

Orthologs of 131 high confidence PfHsp90-dependent proteins for *H. sapiens* (human) and *S. cerevisiae* S288C were identified via Database for Annotation, Visualization, and Integrated Discovery (DAVID) Ortholog tool.^115,116^ All mapped orthologs were considered for each protein. Physical and genetic interactions with HSP90AA1 and HSP90AB1 for human, and HSP82 and HSC82 for yeast were identified with BioGRID 5.0.^117^ Overlap between human and yeast datasets was identified by mapping human Hsp90 interactors to yeast orthologs and identifying shared Hsp90 interactors. To identify proteins unique to Apicomplexa phylum and subsequently, *Plasmodium*, the 131 PfHsp90-dependent proteins were searched in OrthoMCL DB (release 6.21, May 7, 2024).

##### Indispensability analysis

Data for mutagenesis index score (MIS), mutagenesis fitness score (MFS), and mutability was obtained from Zhang *et al*.^58^ The 131 PfHsp90-dependent proteins were assessed based on these parameters, where proteins were classified as essential for *Plasmodium* survival if MIS<0.5, MFS<0 and the gene was non-mutable in coding sequence (CDS).

##### Functional analysis

Gene Ontology (GO) Biological process (BP) and Cellular component (CC) functional enrichment analyses were performed in PlasmoDB^118^ with *p*-value cutoff of 0.05. The enrichment analysis was performed with *P. falciparum* 3D7 selected. The protein-protein interaction network was generated in String version 12.0 with clusters created via k-means clustering and number of clusters set to 4. To identify functional associations of structural homologs of the proteins excluded in GO BP, their AlphaFold files were submitted to the Dali server. Matches against PDB25 were selected and the top 5 structurally similar proteins were submitted to STRING^119^ to identify their GO BP terms. Proteins functionally associated with DNA processes were highlighted. To predict the subcellular localization, proteins excluded by GO CC were searched via the sequence-based prediction tool DeepLoc-2.0 using high-quality model.^63^

##### Enrichment for IDRs and DBRs

IUPRED and ANCHOR scores, which estimate the probability that a residue lies within a disordered region or disordered binding region, were generated for each protein sequence via IUPRED2.^95,120^ Residues with scores greater than 0.5 were classified as positive for IDRs or DBRs. Along with the PfHsp90-dependent proteins, 131 randomly selected *P. falciparum* proteins were analyzed as a reference. The percent of IDR and DBR was quantified for each protein.

#### Thermal Shift Western blot Analysis

Due to the lack of antibodies specific to the cytosolic isoform of PfHsp90 (UniProt ID: Q8IC05), an antibody against human Hsp90 was employed. For validation of antibody specificity, untreated parasites were released from host RBCs at 10-15% parasitemia and 1% hematocrit at the trophozoite stage (22-38 hpi). The pellets were then reconstituted in 4 volumes of buffer (PBS supplemented with a cOmplete tablet). Separately, RBCs were diluted 1:5 in buffer. RBC and parasite pellets were lysed and clarified as described above. Parasite lysate was subsequently diluted to 1.5 mg/mL, while lysed RBCs were diluted 1:100 in buffer. For inhibitor and heat treatments, *P. falciparum* lysates were diluted to 1.5 mg/mL, split into aliquots of equal volume, and incubated at RT for 1 hr with 100 µM GA, 100 µM XL, 100 µM fosmidomycin at a final DMSO or PBS concentration of 5%. Following drug treatment, the sample was split into equal aliquots and subjected to a 3 min heat treatment via a C1000 Touch PCR thermal cycler (Bio-Rad). A temperature range of 47.1 – 53.1 °C was used to identify an appropriate thermal shift window, which indicated a Hsp90 T_m_ shift at 41.9 °C. For band intensity quantification experiments, samples were heated to 49.1 °C in the presence and absence of inhibitors, cooled to RT for 3 min and then placed on ice. Proteins were precipitated via ultracentrifugation at 48,000 RPM at 4 °C for 20 min, after which the soluble fractions were collected and used for the downstream analysis. The samples were resolved on 4-10% Tris glycine gel and transferred to a nitrocellulose membrane via Mini Trans-Blot cell (Bio-Rad). The membrane was stained with Ponceau S (Sigma) for 15 min, rinsed and imaged. The stain was subsequently removed by washing the membrane with 0.1 M NaOH for 1 min, after which the membrane was rinsed and blocked with 3% BSA in PBST (PBS with 0.2% Tween 20 (Sigma)). Following three washes with PBST, the membrane was incubated with monoclonal anti-Hsp90 antibody (Sigma) at 1:5,000 for 1 hr at RT.

After three PBST washes, the membrane was probed with Alexa Fluor 488-conjugated donkey anti-mouse IgG (H+L) secondary antibody (ThermoFisher Scientific) at 1:1,000 for 1 hr at RT. Following PBST washes, the fluorescence signal was measured via a ChemiDoc MP System (Bio-Rad). Analysis and band quantification was performed in ImageLab 6.1, where band intensities of heat-treated samples were normalized to their respective RT controls. Experiments were performed in four biological replicates, with technical duplicates of DMSO.

#### Fixed fluorescence and immunofluorescence microscopy staining

##### Blood stage

*P. falciparum* parasites at 30 hpi were added to a 12-well tissue culture plate (GenClone) and supplemented with 10 µM of 7 EdA (Biosynth), as well as GA (0.3 µM and 3 µM), XL (0.3 µM and 3 µM), pyrimethamine (0.05 µM and 0.5 µM), or fosmidomycin (1.17 µM and 11.7 µM). All compounds were dissolved in DMSO except for fosmidomcyin, which was dissolved in PBS. For control experiments, parasites were similarly supplemented with 10 µM EdA and treated with cipargamin (0.0004 µM), chloroquine (0.02 µM), dihydroartemisinin (0.0004 µM) and XL888 (0.3 µM). The final DMSO concentration was 0.5% in a final volume of 1 mL for all conditions. Parasites were incubated for 12 hrs at 37 °C with 5% CO_2_. Following incubation, 500 µL from each well was centrifuged, washed with 1 mL of PBS, and then fixed with 500 µL of fresh 4% (wt/vol) paraformaldehyde (Sigma) and 0.0075% (vol/vol) glutaraldehyde (Sigma) in PBS for 30 min in the dark at RT. The cells were subsequently washed with PBS, and permeabilized in 500 µL of 0.1% (vol/vol) TritonX-100 in PBS for 10 min at RT in the dark. Following 2 washes with PBS, cells were blocked in freshly prepared, filter-sterilized 3% (wt/vol) bovine serum albumin (Equitech Bio) and 300 mM glycine (VWR International) in PBS at 4 °C overnight with rotation. Cells were washed twice with PBS and then stained with a freshly prepared solution containing 2 mM CuSO_4_ (ChemCruz), 12 µM Azide-fluor 488 (Sigma), and 10 mM sodium ascorbate (Sigma) in PBS for 1 hr in the dark at RT. Following this, cells were washed three times with PBS and stained with 10 µg/mL Hoechst 33342 (ThermoFisher Scientific) in PBS for 15 min in the dark at RT. Cell pellets were then washed three times with PBS, reconstituted in 50 µL of PBS and stored at 4 °C in the dark prior to imaging. Cells were added to a 384-well glass bottom plate (Cellvis) and diluted in PBS to obtain a monolayer. The plate was briefly centrifuged and imaged using a Leica MICA inverted confocal microscope. A 20x 0.75 NA DRY objective was used for quantification, while 63x 1.40 NA OIL objective was used for representative images. Laser illumination was with diodes for 405 nm and 488 nm. Images were acquired sequentially by line scanning bidirectionally with line averaging of 4 and pinhole set to 1 Airy unit via Leica LASX software (version 1.4.7) for quantification, and the line averaging of 2 for representative images.

##### Liver stage

HepG2 cells (5,000 cells/well) were seeded on glass 384-well plates (Cellvis-P38415HN) and allowed to recover overnight before infection with GFP-expressing *P. berghei* sporozoites (4,000/well). At 24 hours post-infection, GA (1 and 10 µM), XL (1 and 10 µM), pyrimethamine (0.025 and 0.25 µM), and DMSO were added to the media at the same time as 5 µM EdA. For control experiments, parasites were similarly treated with cyclosporin A (0.0085 and 0.085 µM), cycloheximide (0.235 and 2.35 µM), and XL (1 and 10 µM) along with 10 µM EdA . At 30 hours post-infection, cells were fixed in 4% paraformaldehyde for 10 min and then washed with PBS. Cells were permeabilized with 0.1% TritonX-100 in PBS for 10 min, washed with PBS, and incubated with blocking buffer (0.05% PBST with 3% BSA) for 1 h. The primary antibody FITC conjugated-goat anti-GFP (abcam cat#AB6662, 1:1000) was diluted in blocking buffer for 1 hr at RT. Cells were washed and the copper catalyzed click reaction was performed as described above using 12 µM AZDye594-Azide (Jena Bioscience) for 1 hr at RT in the dark. After antibody staining, cells were washed in PBST and nuclei were stained with DAPI (0.5 μg/mL) for 10 min at RT. Cells were washed with PBS before imaging using a Leica MICA inverted confocal microscope. A 20x 0.75 NA DRY objective was used for quantification, while 63x 1.40 NA OIL objective was used for representative images. Laser illumination was with diodes for 405, 488, and 561 nm. Images were acquired sequentially by line scanning bidirectionally with line averaging of 4 and pinhole set to 1 Airy unit via Leica LASX software (version 1.4.7) for quantification, and the line averaging of 2 for representative blood stage and 8 for representative liver stage images.

#### Microscopy image analysis

Image analysis was performed in FIJI. Regions of interest were identified via Otsu auto threshold in the Hoechst channel for blood stage and GFP channel for liver stage parasites. To ensure complete capture of segmented schizont nuclei per infected cell, the following morphological operations were applied in the following order: “Dilate”, “Close-”, “Fill holes”. The resulting ROIs were subsequently applied to EdA channel. Integrated density of the EdA channel was measured for blood stage. In contrast to RBCs, *Plasmodium*-infected hepatocytes can vastly vary in size throughout the infection. Hence, to ensure that the EdA incorporation assessment was not impacted by changes in parasite size, the mean gray value of EdA was quantified for the liver stage. To ensure equal sample size across conditions for the blood stage analysis, a randomly selected subset of 300 parasites from each treatment was used for statistical analysis and data representation. Parasites with integrated density of >1500 and MGV >8 were visually inspected, subsequently classified as artifacts and excluded from the analysis. Samples were analyzed across three biological replicates.

#### Quantification and statistical analysis

GraphPad Prism 10 software was employed for data analysis. The sample size, standard error, and statistical tests used in analysis are noted in respective figure legends.

## AKNOWLEDGEMENTS

We would like to acknowledge the Duke University School of Medicine Proteomics and Metabolomics Core Facility, especially Drs. Erik Soderblom, Jeffrey Kuhn, and Matt Foster for their valuable assistance in acquiring the LC-MS/MS data. We are grateful to Dr. Christopher R. Mansfield for insightful feedback, discussions, and for providing purified HsHsp90. We also thank Dr. Anna Truong for feedback and support with molecular docking efforts. Finally, we are grateful to the members of the Derbyshire lab for their helpful feedback and review of the manuscript. Schematics were created using BioRender. This research was supported by the National Institute of Health (R01 Al173295), the Sloan Research Foundation, and a Camille Dreyfus-Teacher Scholar Award to E.R.D. Fellowship support was provided to G.I. through Bradsher fellowship and to C.J.H. through the Amgen Scholar Program.

## AUTHOR CONTRIBUTIONS

G.I. and E.R.D. conceptualized the study. G.I. performed the blood stage inhibition and TSA assays, co-IP, ortholog analysis, cell staining for blood stage parasites, and image acquisition for blood and liver stages. Y.C. prepared samples for the LC-MS/MS experiments and helped analyze the LC-MS/MS data. G.I., Y.C., M.C.F. and E.R.D. interpreted the LC-MS/MS results. M.E.C. performed liver stage infections and cell staining. G.I. and C.J.H. performed protein expression and purification, and bioinformatic and microscopy image analyses. G.I. and E.R.D. wrote the manuscript and designed the experiments. All authors reviewed and provided feedback on the manuscript.

## DECLARATION OF INTERESTS

The authors declare no competing interests.

## DECLARATION OF GENERATIVE AI AND AI-ASSISTED TECHNOLOGIES IN THE WRITING PROCESS

During the preparation of this work, the authors used ChatGPT and Perplexity in order to improve grammar, language, and readability. After using these tools, the authors reviewed and edited the content as needed and take full responsibility for the content of the published article.

## Supplemental information

Document S1. Figures S1-S14

Table S1. Proteins identified in global proteomic analysis, related to Figure 1.

Table S2A-F. PfHsp90-dependent proteins and associated analyses, related to Figures 1 and 2

Table S3A-D. Functional analyses of PfHsp90-dependent proteins, related to Figures S2 and S14.

## Notes

### Competing Interest Statement

The authors have declared no competing interest.

