## Supplementary material for "Chemoproteomic profiling of *Plasmodium falciparum* Hsp90 inhibition reveals functional link to DNA replication pathways": Document S1

**SUPPLEMENTAL:**

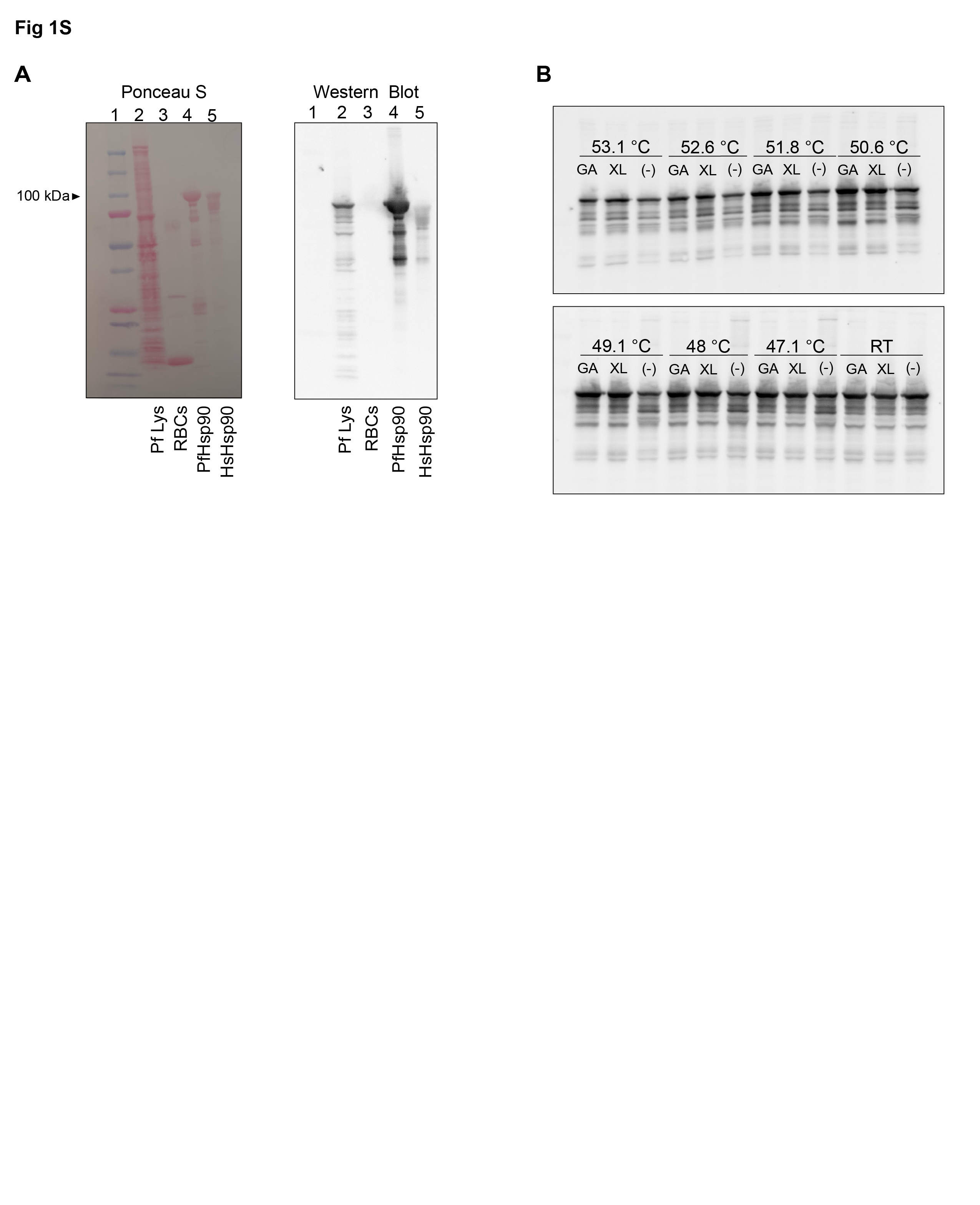

**Fig S1**. **Hsp90 antibody specificity and T_m_ determination.** (A) Representative Ponceau S total protein stain (left panel) and Western blot (right panel) probed with anti-Hsp90 antibody (expected size ~100 kDa). Lane 1, Bio-Rad Precision Plus Protein Dual Color Standards; Lane 2, lysate of *P. falciparum* released from infected RBCs (PfLys); Lane 3, lysate of uninfected RBCs; Lane 4, purified PfHsp90; Lane 5, purified HsHsp90. Imaging confirms PfHsp90 detection in parasite lysate but not uninfected RBCs. *n*=3 biological replicates. (B) *P. falciparum* lysate treated with 100 µM of GA or XL and subjected to a 47.1 – 53.1 °C thermal challenge or room temperature (RT). The vehicle control was 5% DMSO (-).

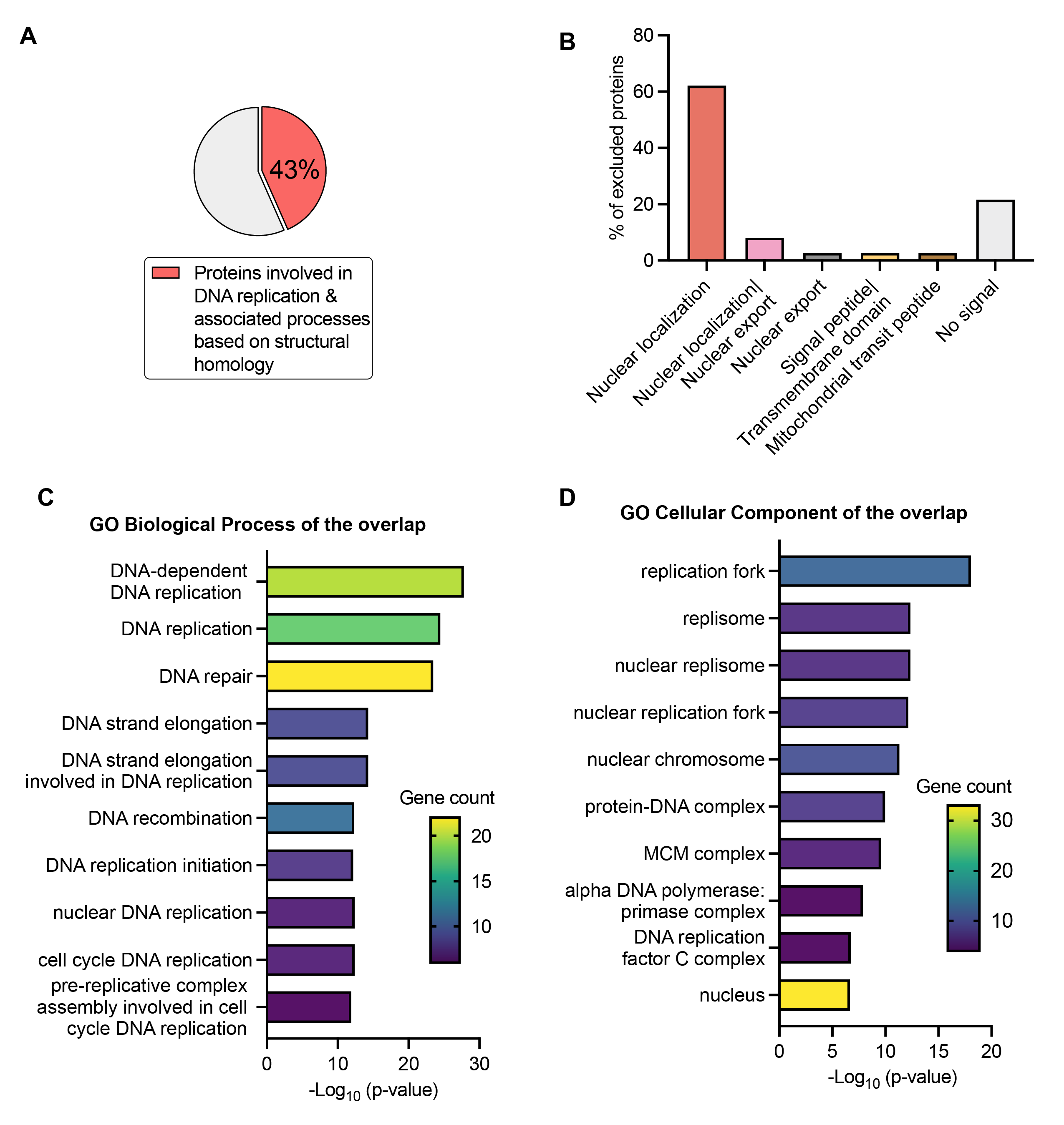

**Fig S2. Computational analysis to inform function of PfHsp90-dependent proteins excluded from GO analyses.** (A) Proteins excluded by GO BP analysis were submitted to Dali server to identify structural homologs, which were then analyzed for their functional associations via STRING. Of the 51 excluded proteins, 22 (43%) had structural homologs involved in DNA replication and associated pathways. (B) Proteins excluded by GO CC analysis were examined via DeepLoc 2.0 to predict their subcellular localization. Of the 37 excluded proteins, 23 (62%) were enriched in nuclear localization motifs, 3 (8%) were enriched in nuclear localization|nuclear export motifs, 1 (3%) was enriched in nuclear export, signal peptide|transmembrane domain, or mitochondrial transit peptide motifs. No signal was detected in 8 (22%).

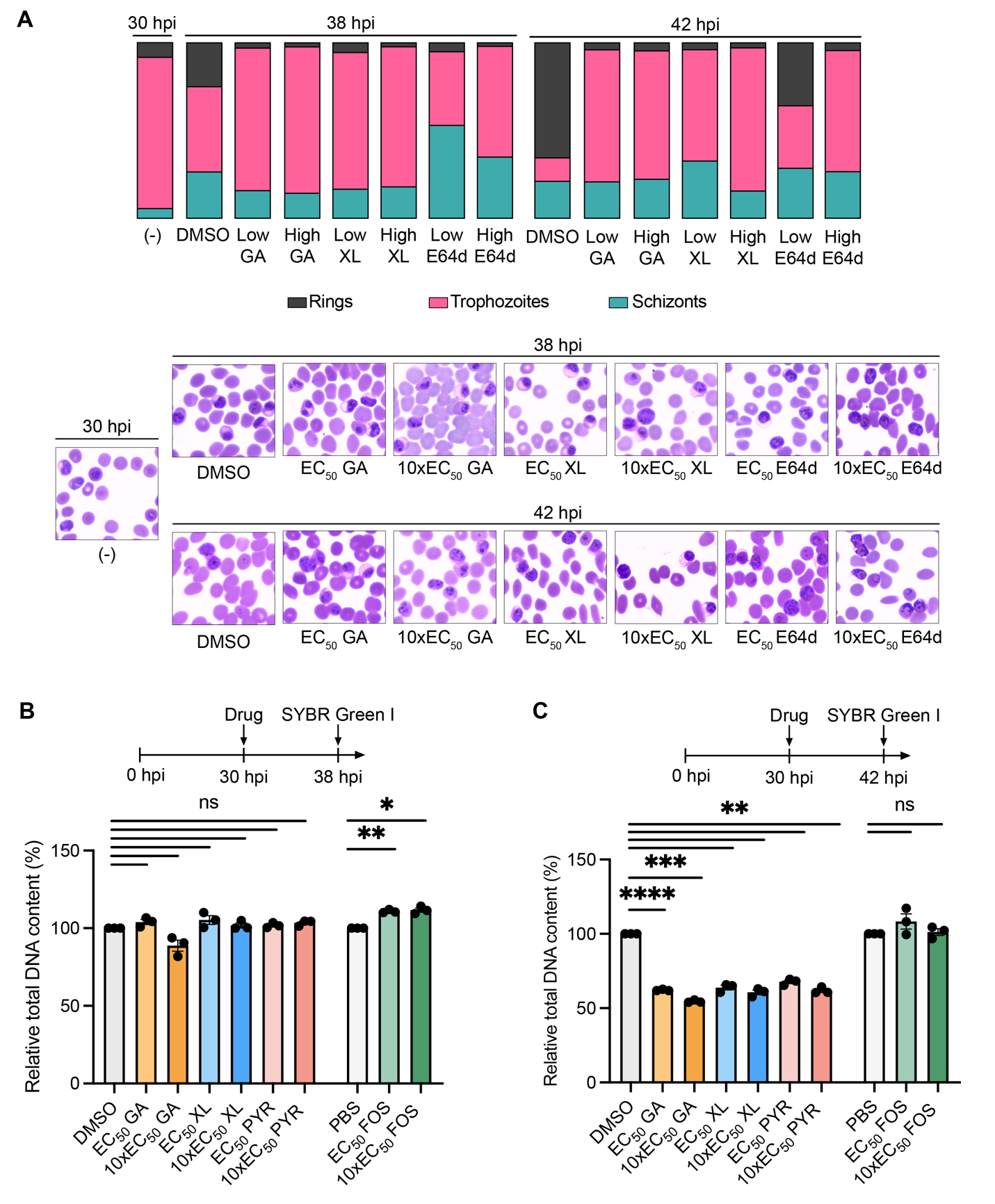

**Fig S3. Impact of PfHsp90 inhibitors administered at EC_50_ and 10xEC_50_ concentrations for 8 or 12 hours on stage progression and DNA quantity.** (A) *P. falciparum*-infected RBCs were treated with GA, XL, and E64d at EC_50_ (low) and 10xEC_50_ (high) concentrations for 8 hours from 30 to 38 hpi or for 12 hours from 30 to 42 hpi. DMSO was at 0.5%. Data for 30 hpi DMSO, and 42 hpi GA, XL, and E64d at EC_50_ is reproduced from **Fig 4A** to represent the complete dataset. Following incubation, stage progression was assessed via light microscopy analysis of smears and the parasites were scored as rings, trophozoites, or schizonts. Data shown as *n*=3 biological replicates, analyzing >290 parasites for each condition. (B and C) *P. falciparum*-infected RBCs were treated with GA, XL, PYR, and FOS at EC_50_ and 10XEC_50_ concentrations for 8 hours from 30 hpi to 38 hpi (B) or for 12 hours from 30 hpi to 42 hpi (C). PYR was the positive control while FOS was used as the inactive negative control. The vehicle controls were at 0.5% (GA, XL, and PYR: DMSO and FOS: PBS). Following incubation, lysis buffer containing 1xSYBR Green dye was added and the fluorescence signals of each treatment condition were normalized to the signals of their respective vehicle controls. Data for DMSO, PBS and compounds at their EC_50_ concentration is reproduced from **Fig 4B** to represent the complete dataset. Data shown as means ± SEM, *n*=3 biological replicates; not significant (ns) > 0.05, *<0.05, **<0.01, ***<0.001, ****<0.0001; Welch’s t-test.

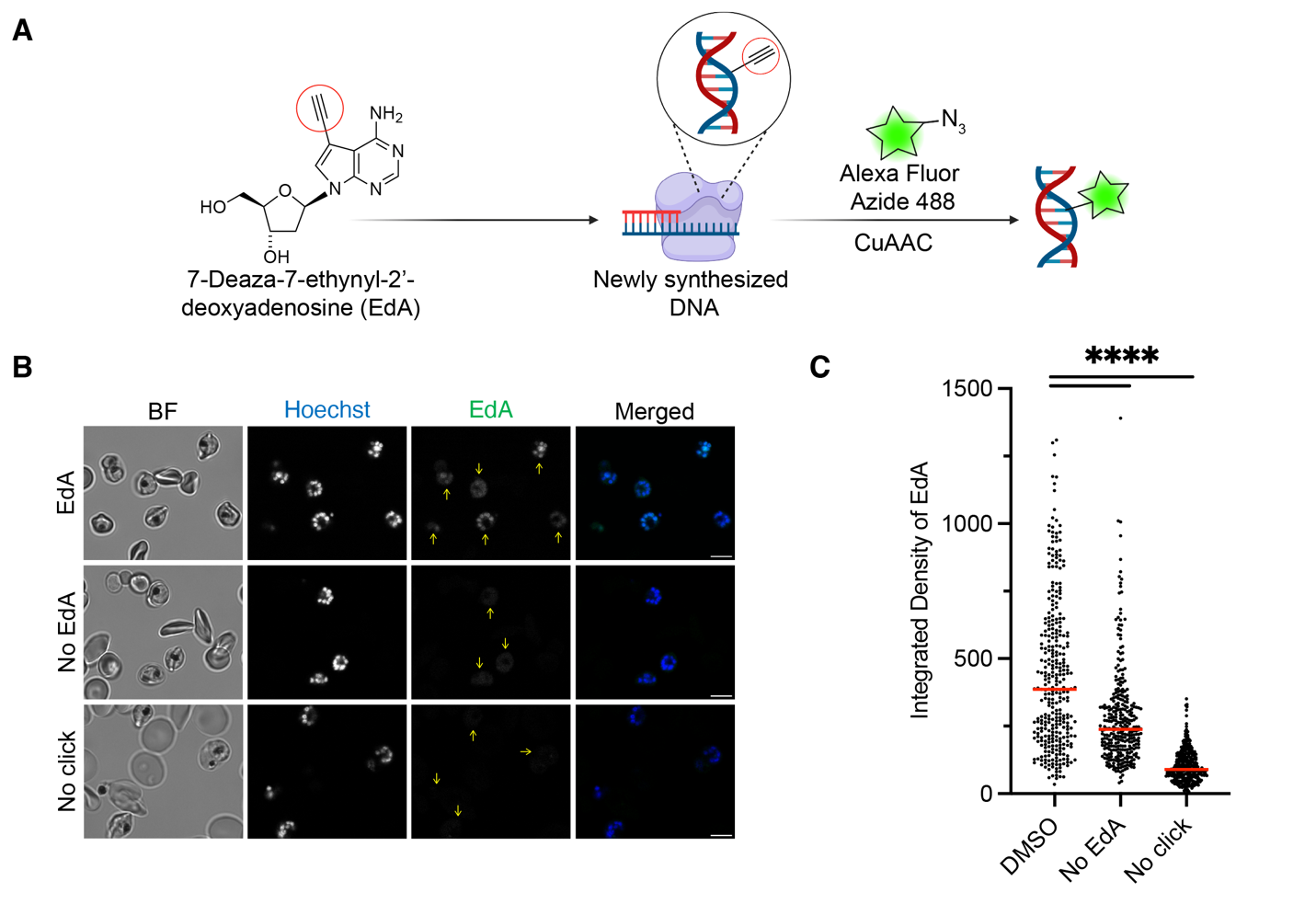

**Fig S4. Validation of EdA incorporation and labeling specificity during the blood stage.** (A) The alkyne-containing nucleoside EdA incorporates into newly synthesized *P. falciparum* DNA, allowing for Cu(I)-catalyzed azide-alkyne cycloaddition (CuAAC) with Alexa Fluor Azide 488. (B-C) *P. falciparum*-infected RBCs incubated with 10 µM EdA, no EdA, or no Alexa Fluor Azide 488 (No click) at 0.5% DMSO for 12 hours at 30 hpi, fixed at 42 hpi and stained. (B) Representative confocal fluorescence microscopy images of *P. falciparum*-infected RBCs stained Alexa Fluor Azide 488 (green) to label newly synthesized DNA and Hoechst (blue) to label nuclei . Yellow arrows highlight infected cells. Scale bars are 5 µm. (C) The integrated density in the EdA channel was quantified per parasite. Data for EdA (DMSO) was reproduced from **Fig 4D**. Median indicated with red bar, *n*=3 biological replicates, analyzing >300 parasites for each condition; ****<0.0001; Mann-Whitney U test.

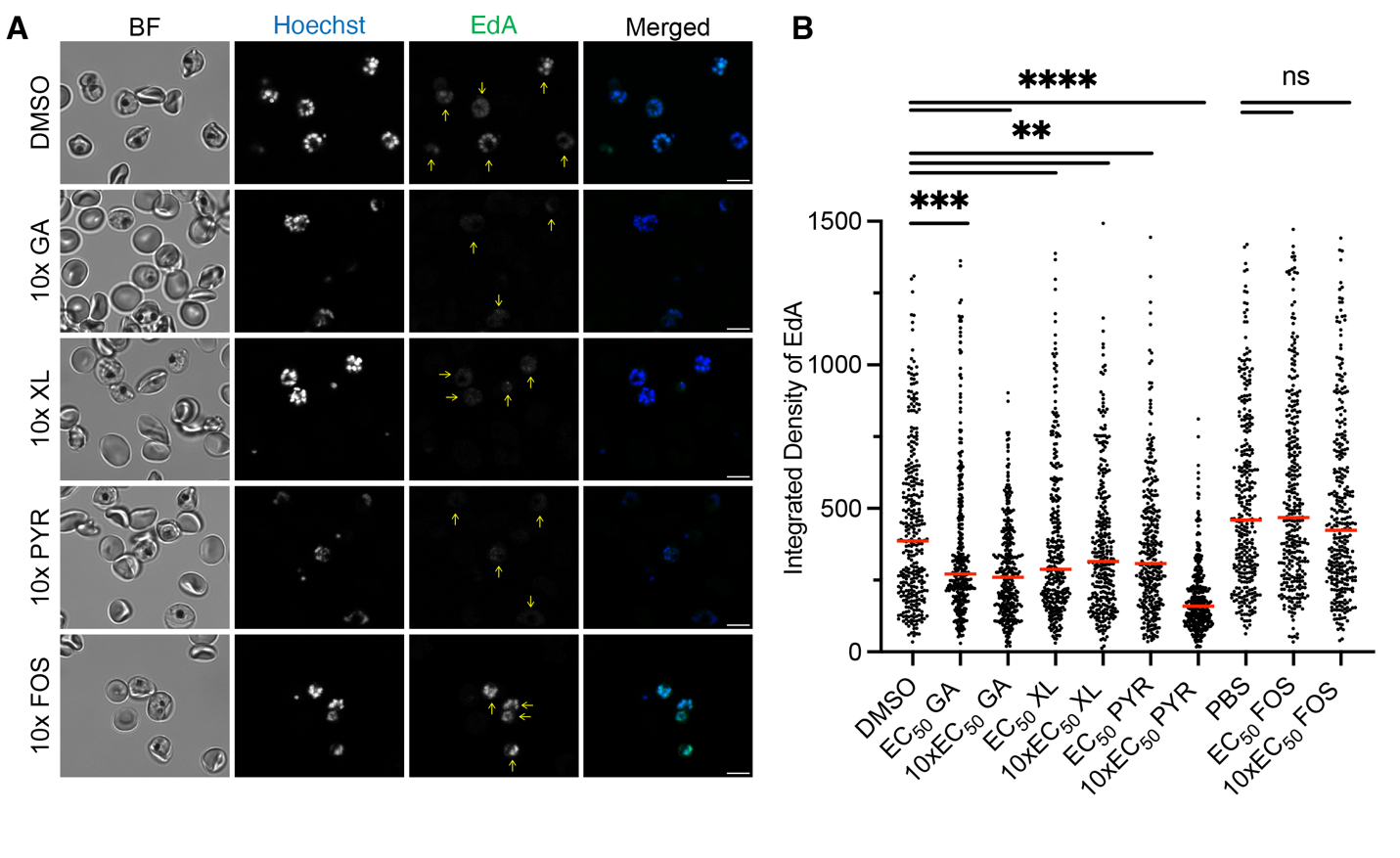

**Fig S5. Impact of PfHsp90 inhibition at EC_50_ and 10xEC_50_ concentrations on DNA synthesis during the blood stage.** (A-B) *P. falciparum*-infected RBCs incubated with 10 µM EdA and drugs at EC_50_ and 10xEC_50_ concentrations for 12 hours at 30 hpi, fixed at 42 hpi and stained. (A) Representative confocal fluorescence microscopy images of *P. falciparum*-infected RBCs stained with Alexa Fluor Azide 488 (green) to label newly synthesized DNA and Hoechst (blue) to label nuclei. Yellow arrows highlight infected cells. Scale bars are 5 µm. (B) The integrated density of EdA was quantified per parasite. Data for DMSO, PBS and compounds at their EC_50_ concentration is reproduced from **Fig 4D** to represent the complete dataset. PYR was the positive control while FOS was used as the inactive negative control. The vehicle controls were at 0.5% (GA, XL, and PYR: DMSO and FOS: PBS). Median indicated with red bar, *n*=3 biological replicates, analyzing >300 parasites for each condition; not significant (ns) > 0.05, **<0.01, ***<0.001, ****<0.0001; Mann-Whitney U test.

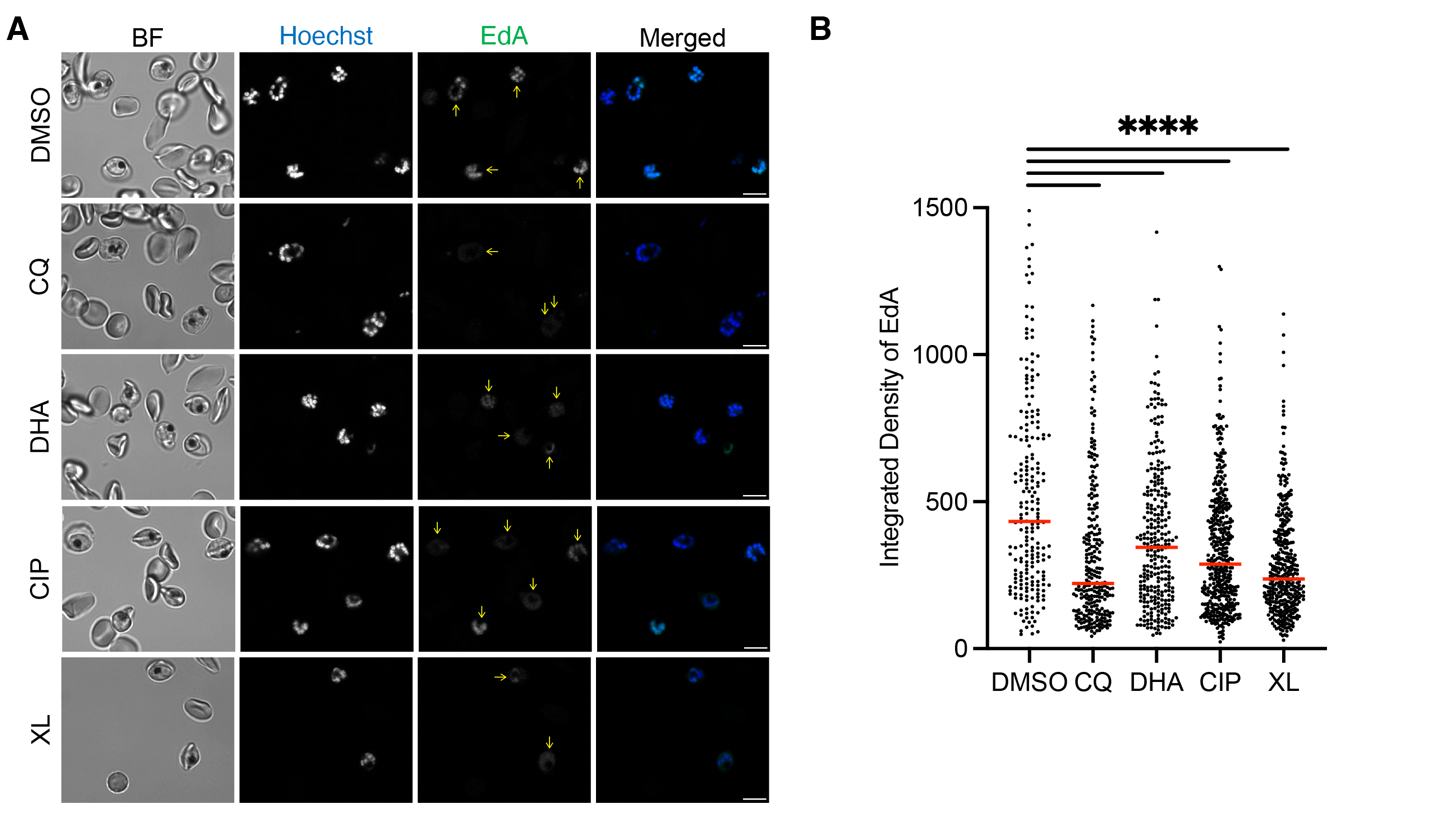

**Fig S6. Impact of treatments with fast-acting antimalarial control compounds on EdA incorporation during the blood stage.** (A-B) *P. falciparum*-infected RBCs were incubated with EdA and drugs at EC_50_ concentrations for 12 hours at 30 hpi, fixed at 42 hpi and stained. (A) Representative confocal fluorescence microscopy images of *P. falciparum*-infected RBCs stained with Alexa Fluor Azide 488 (green) to label newly synthesized DNA and Hoechst (blue) to label nuclei. Yellow arrows highlight infected cells. Scale bars are 5 µm. (B) The integrated density of EdA was quantified per parasite. XL was used as the reference treatment. DMSO was at 0.5%. Median indicated with red bar, *n*=3 biological replicates, analyzing >210 parasites for each condition; ****<0.0001; Mann-Whitney U test.

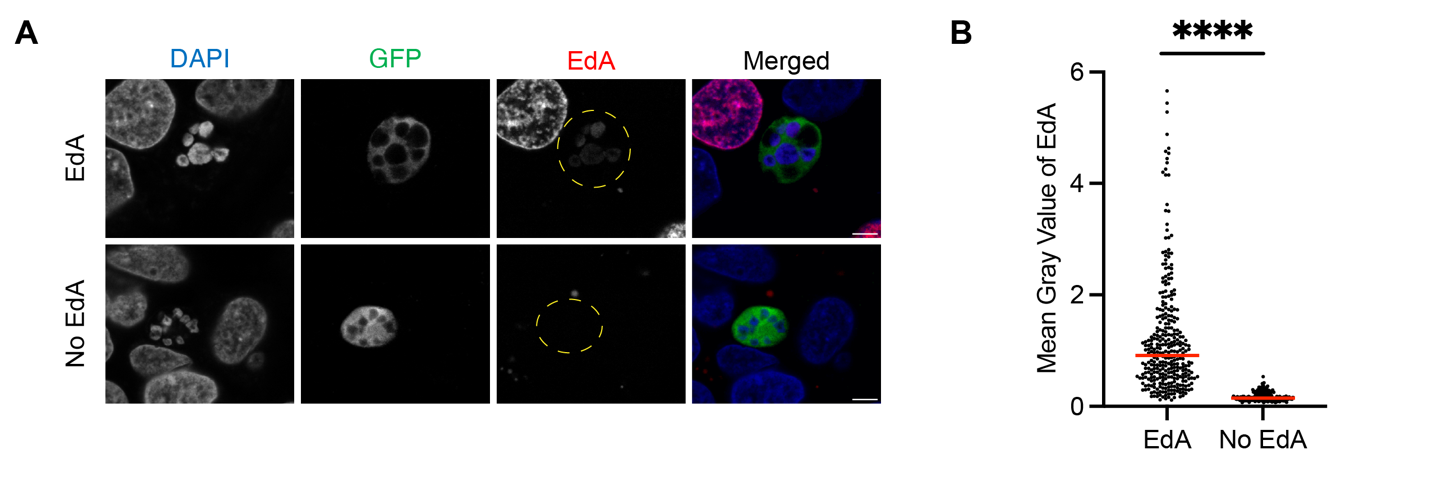

**Fig S7. Validation of EdA incorporation during the *P. berghei* liver stage.** (A-B) HepG2 cells were infected with GFP-expressing *P. berghei*, treated with or without 5 µM EdA for 6 hours at 24 hpi, fixed at 30 hpi and stained. (A) Representative confocal immunofluorescence microscopy images of *P. berghei*-infected cells stained with anti-GFP (green) to label the parasites, AZDye594-Azide (red) to label newly synthesized DNA, and DAPI (blue) to label nuclei. Yellow dashed circles highlight infected cells. Scale bars are 5 µM. (B) The mean gray value of the EdA channel of individual *P. berghei*-infected HepG2 cells was quantified. Data for EdA (DMSO) was reproduced from **Fig 5B**. Median indicated with red bar, *n*=3 biological replicates analyzing >250 parasites for each condition; ****<0.0001; Mann-Whitney U test.

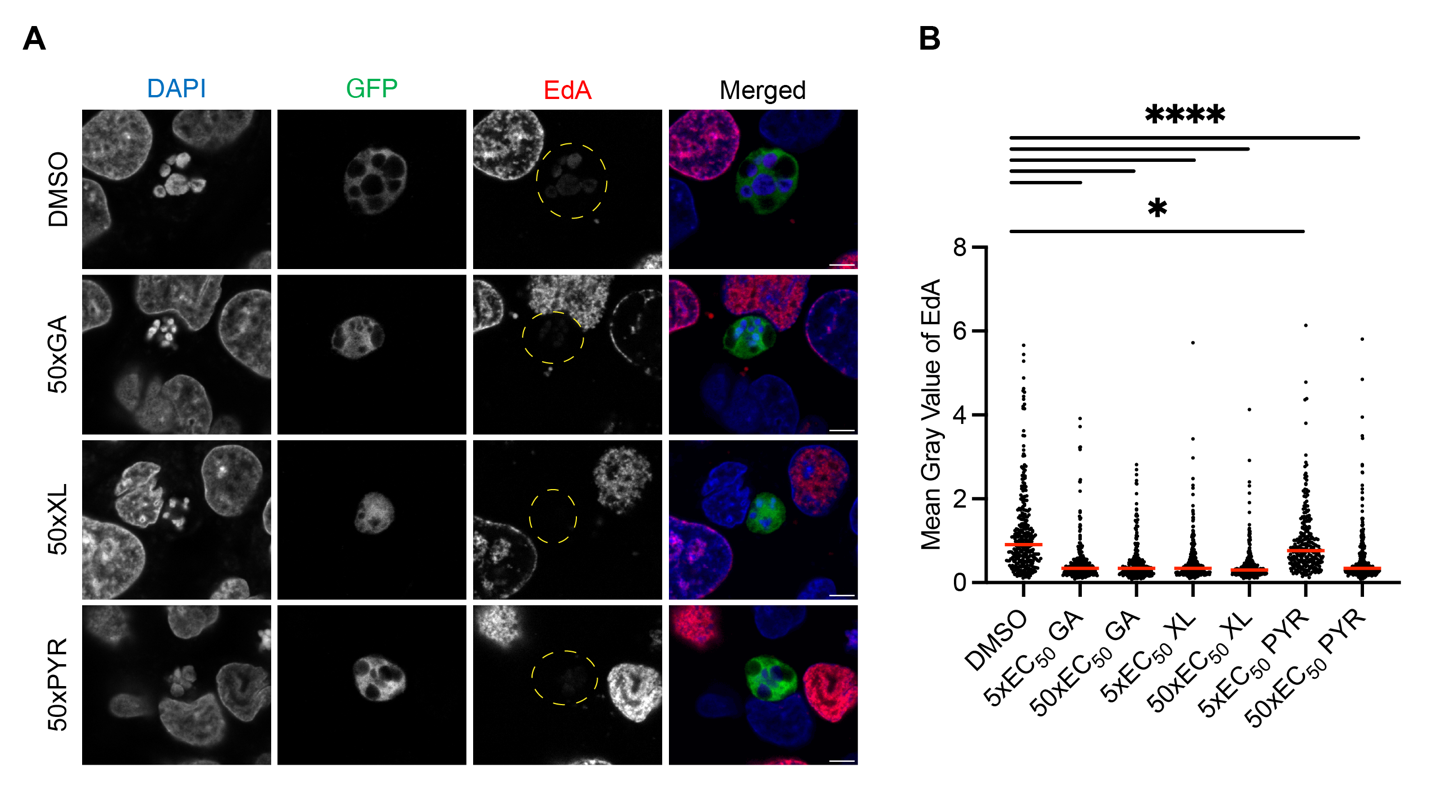

**Fig S8. Impact of Hsp90 inhibition at 5xEC_50_ and 50xEC_50_ concentrations on EdA incorporation during the *P. berghei* liver stage.** (A-B) HepG2 cells were infected with GFP-expressing *P. berghei*, treated with EdA and drugs at 5xEC_50_ and 50xEC_50_ concentrations for 6 hours at 24 hpi, fixed at 30 hpi and stained. (A) Representative confocal immunofluorescence microscopy images of *P. berghei*-infected cells stained with anti-GFP (green) to label the parasites, AZDye594-Azide (red) to label newly synthesized DNA, and DAPI (blue) to label nuclei. Yellow dashed circles highlight infected cells. Scale bars are 5 µm. (B) The mean gray value of the EdA channel of individual *P. berghei*-infected HepG2 cells was quantified. PYR was the positive control. Data for DMSO and compounds at 5xEC_50_ concentration is reproduced from **Fig 5B** to represent the complete dataset. Median indicated with red bar, *n*=3 biological replicates, analyzing >250 parasites for each condition; *<0.05, ****<0.0001; Mann-Whitney U test.

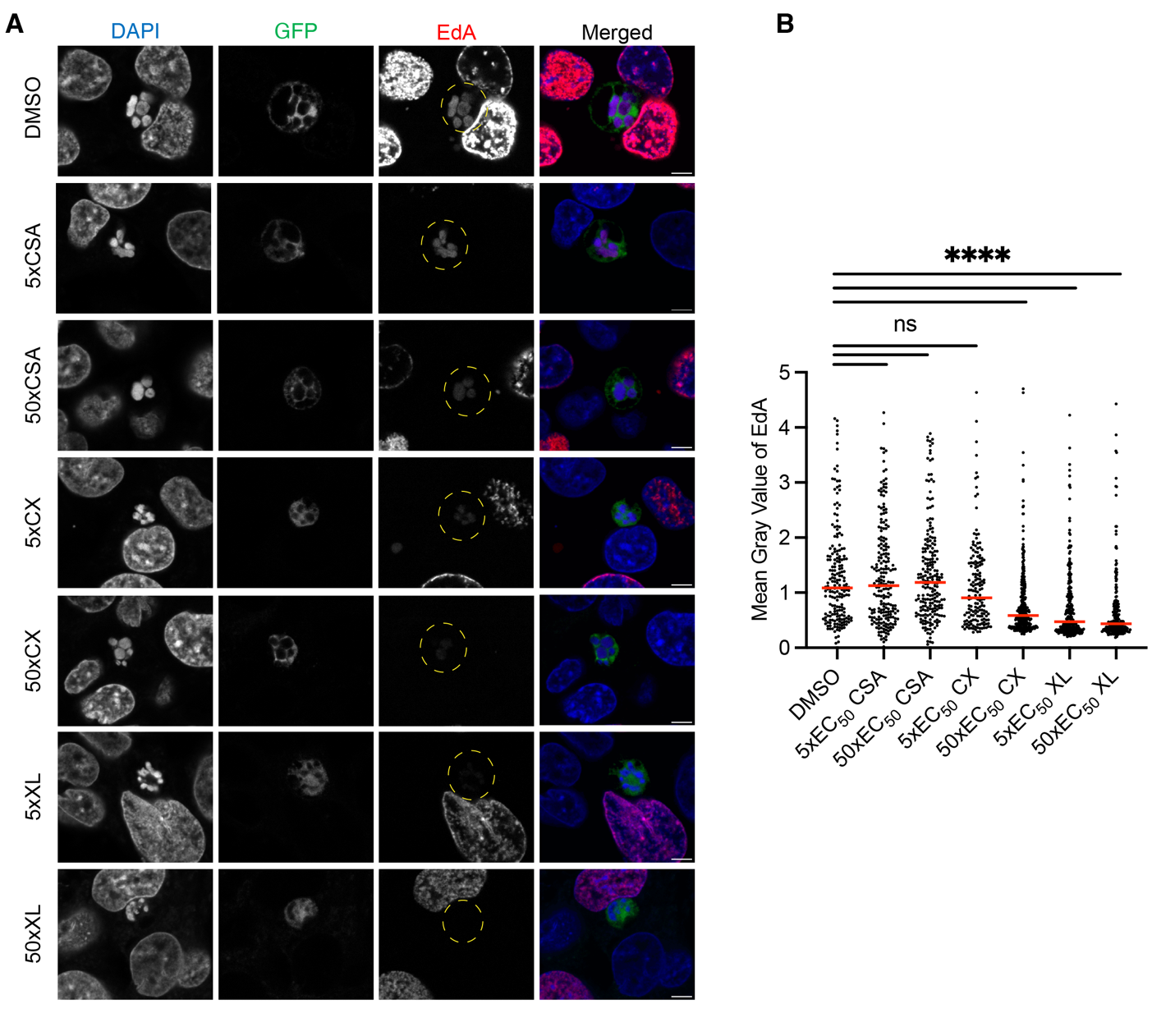

**Fig S9. Impact of treatments with inactive and active control compounds on EdA incorporation during the *P. berghei* liver stage.** (A-B) HepG2 cells were infected with GFP-expressing *P. berghei*, treated with EdA and drugs at 5xEC_50_ and 50xEC_50_ concentrations for 6 hours at 24 hpi, fixed at 30 hpi and stained. (A) Representative confocal immunofluorescence microscopy images of *P. berghei*-infected HepG2 cells stained with anti-GFP (green) to label the parasites, AZDye594-Azide (red) to label newly synthesized DNA, and DAPI (blue) to label nuclei. Yellow dashed circles highlight infected cells. Scale bars are 5 µm. (B) The mean gray value of the EdA channel of individual *P. berghei*-infected HepG2 cells was quantified. Median indicated with red bar, *n*=3 biological replicates, analyzing >150 parasites for each condition; not significant (ns) >0.05, ****<0.0001; Mann-Whitney U test.

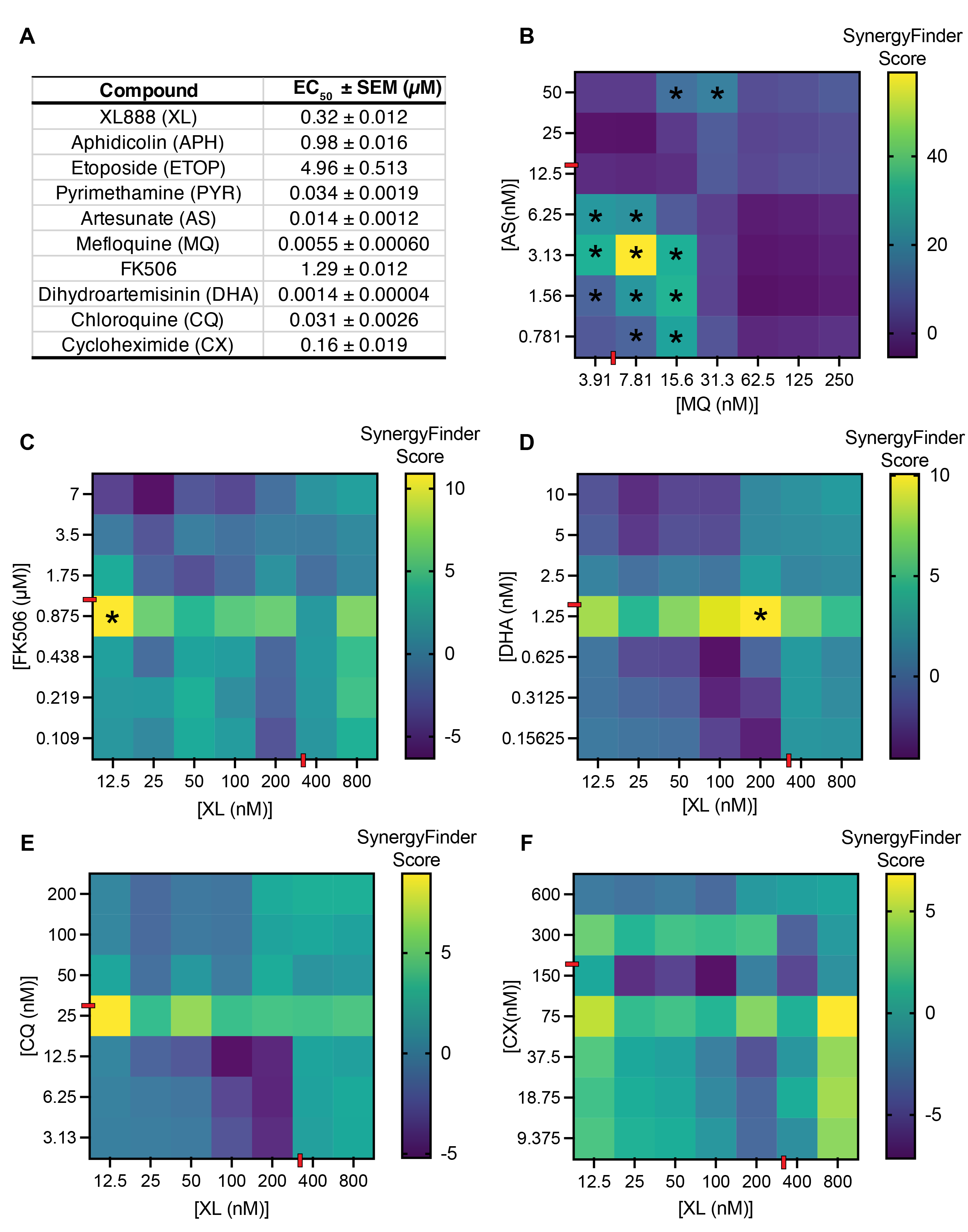

**Fig S10. Synergy analysis of mechanistically diverse compounds with XL and established controls.** (A) Table of EC_50_ values generated from dose response curves of XL, APH, ETOP, PYR, AS, MQ, FK506, DHA, CQ and CX inhibition of *P. falciparum* 3D7 parasites from the synergy assays. Data shown as means ± SEM, *n*=3 biological replicates. Determined EC_50_ values for XL, APH, PYR, AS, MQ, FK506, DHA, and CX agree with literature reports,^13,67,79,87,121–124^ while EC_50_ value for ETOP has not been previously reported. (B-F) Synergy heatmaps depicting combined effects of AS with MQ (B), as well as XL with FK506 (C), DHA (D), CQ (E), and CX (F). Synergy was evaluated via SynergyFinder 3.0 zero interaction potency (ZIP) model with baseline correction. Synergistic interactions (SynergyFinder score>10) are marked with an asterisk (*). The red dashes indicate individual EC_50_ concentrations for each compound. Data shown as means, *n*=3 biological replicates.

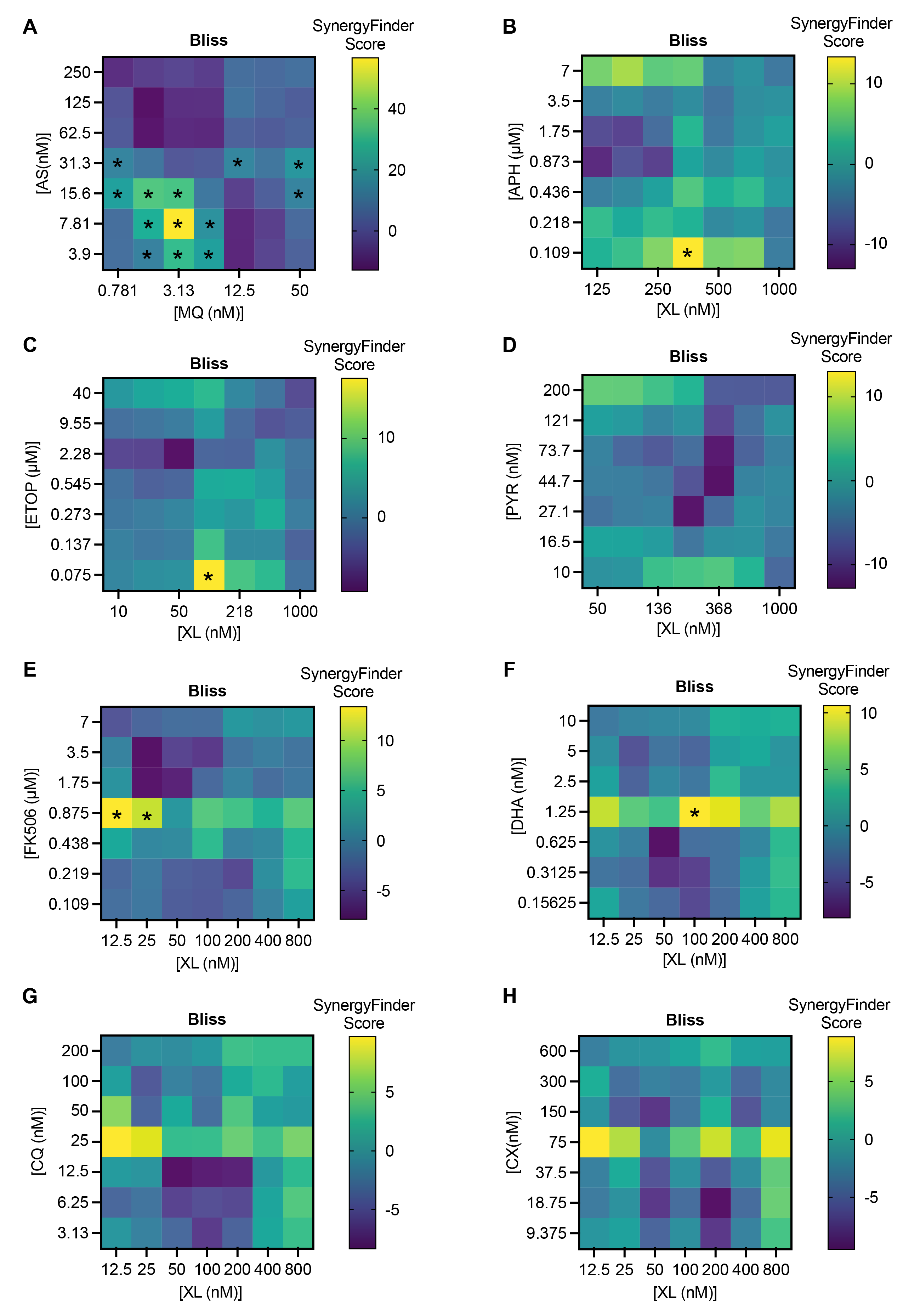

**Fig S11. Synergy analysis via Bliss reference model.** Synergy heatmaps showing combined effects of AS with MQ (A), as well as XL and APH (B), ETOP (C), PYR (D), FK506 (E), DHA (F), CQ (G) and CX (H). Synergy was evaluated via Bliss model with baseline correction. Synergistic interactions (SynergyFinder score>10) are marked with an asterisk (*). Data shown as means, *n*=3 biological replicates.

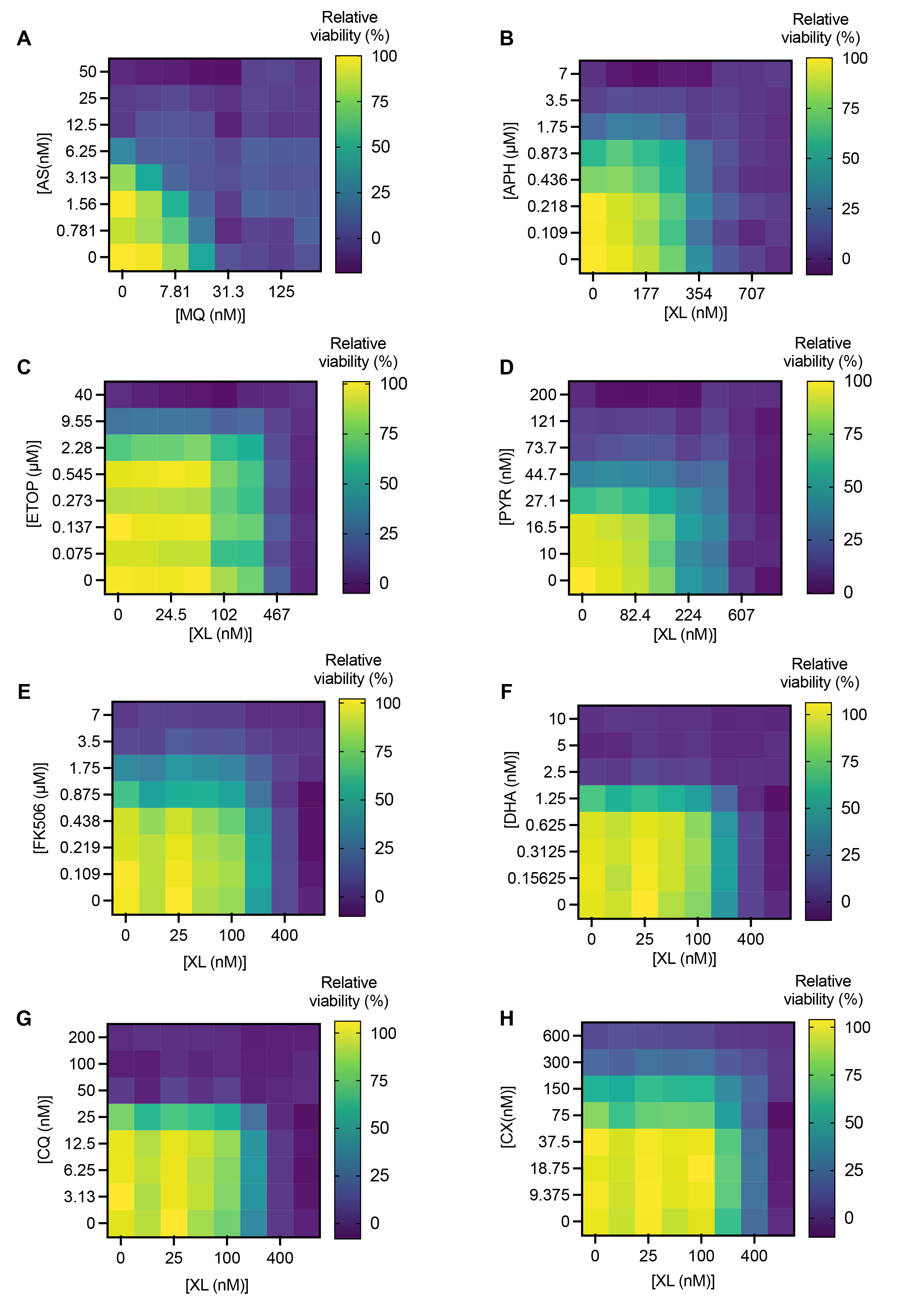

**Fig S12. Assessment of relative parasite viability with compound co-treatments**. Relative parasite viability after co-treatment with AS with MQ (A), as well as XL and APH (B), ETOP (C), PYR (D), FK506 (E), DHA (F), CQ (G) and CX (H). Assessment was via the SYBR Green I assay. Data shown as means, *n*=3 biological replicates.

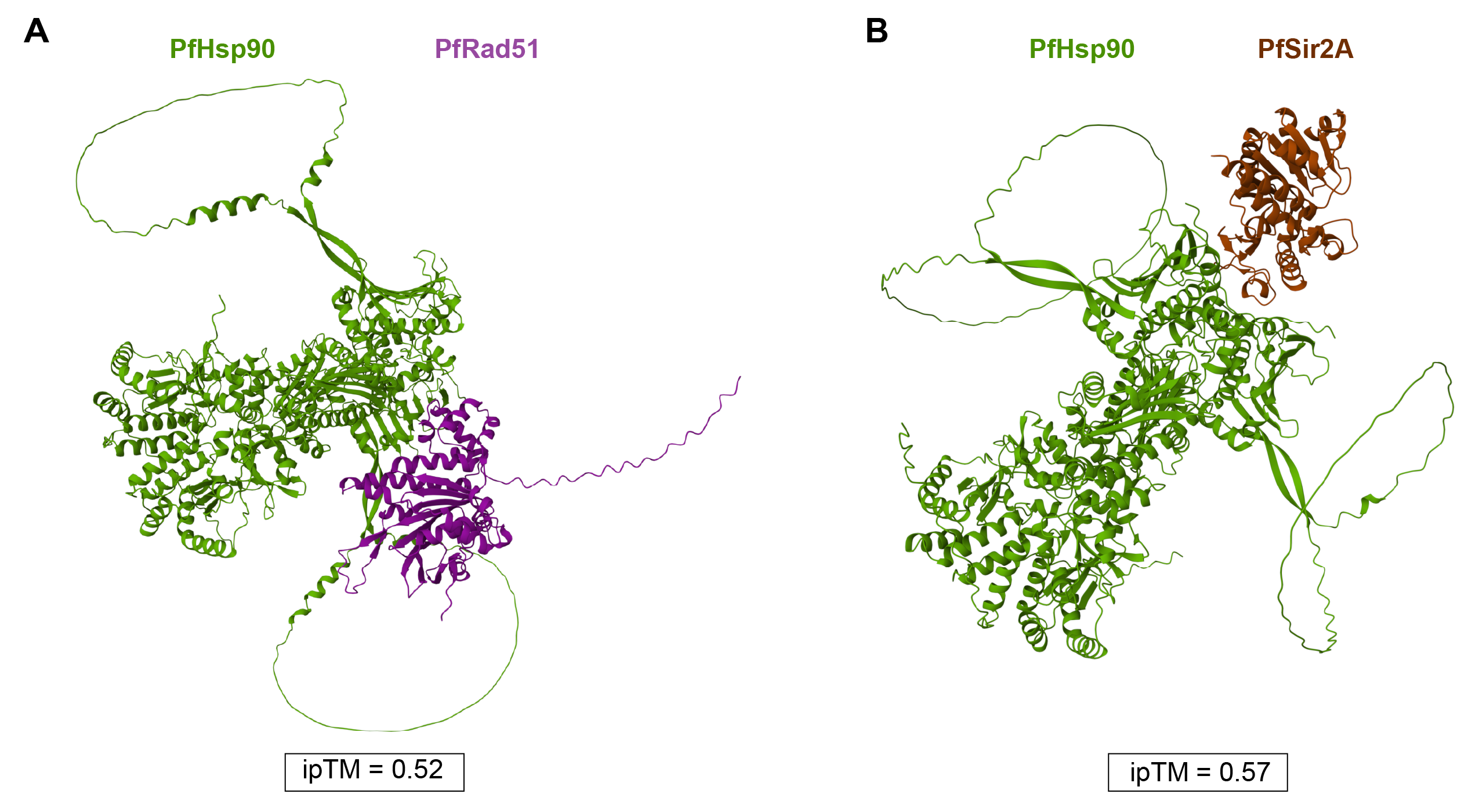

**Fig S13. Prediction of PfHsp90 interactions with known client proteins by AlphaFold 3.** Predictions of interactions between full-length PfHsp90 (green) and PfRad51 (purple) (A) and PfHsp90 and PfSir2A (brown) (B). The accuracy of structural prediction was assessed in AlphaFold 3 via interface predicted template modeling (ipTM) score, where predictions of interactions are classified as high confidence (ipTM>0.8), moderate confidence (between 0.6 and 0.8), and failed prediction (<0.6).

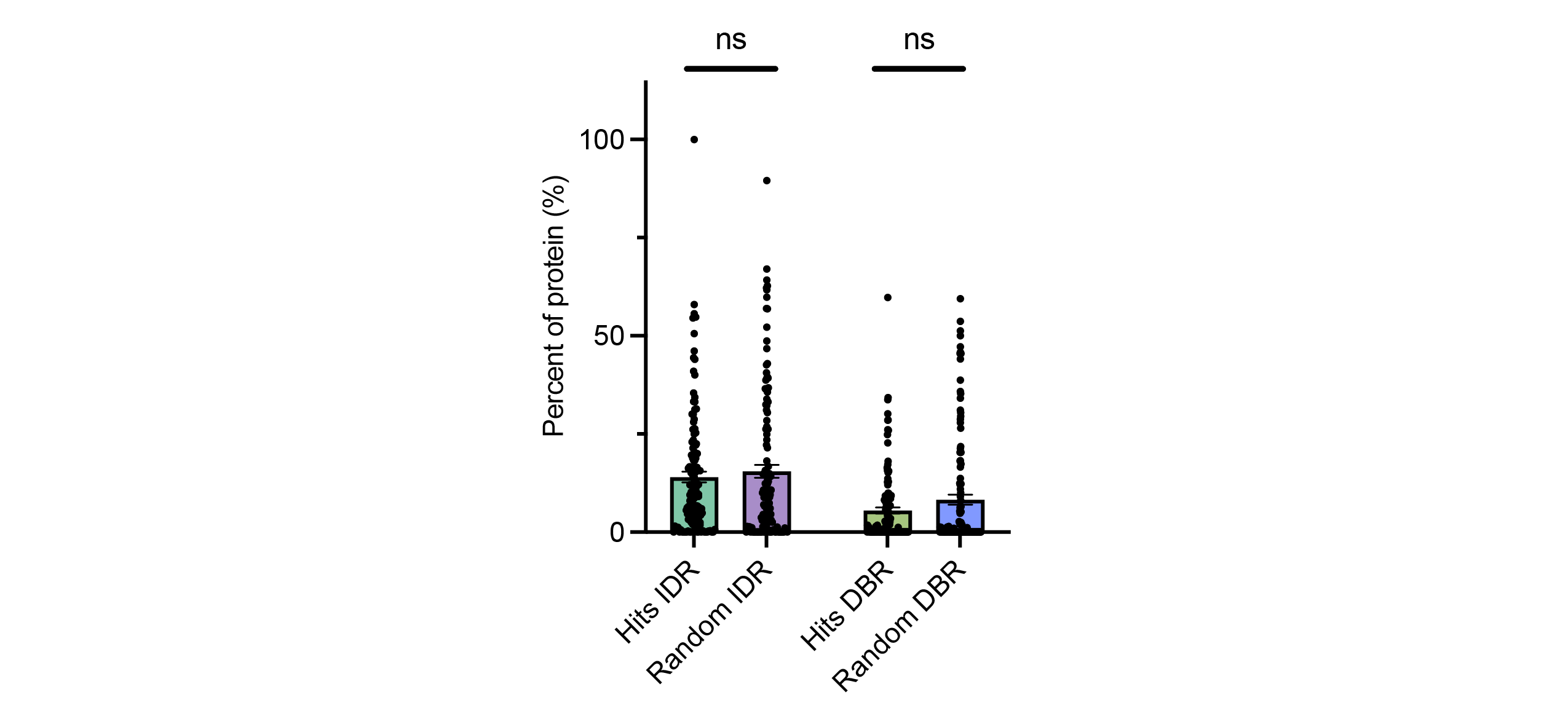

**Fig S14. Assessment of PfHsp90-dependent proteins for IDR and DBR enrichment.** The 131 high confidence PfHsp90-dependent proteins and 131 randomly pooled proteins from the *P. falciparum* proteome were evaluated for percent of intrinsically disordered regions (IDRs) and disordered binding regions (DBRs) per protein sequence. Percent of IDRs and DBRs was quantified via IUPRED2A with a 0.5 cutoff for IUPred2 and ANCHOR2 scores, respectively. Data shown as means ± SEM; not significant (ns) > 0.05; Welch’s t-test.
